# NCS1 regulates Ca^2+^-Dependent Focal Exocytosis of Golgi-derived Vesicles to Help Phagocytic uptake in Macrophages

**DOI:** 10.1101/054858

**Authors:** Nimi Vashi, Syed Bilal Ahmad Andrabi, Swapnil Ghanwat, Mrutyunjay Suar, Dhiraj Kumar

## Abstract

During phagocytic uptake by macrophages, role of Golgi apparatus was previously ruled out. Notably all such reports were limited to Fcγ-receptor mediated phagocytosis. Here we unravel a highly devolved mechanism for recruitment of Golgi-derived secretory vesicles during phagosome biogenesis, which was important for uptake of most cargos except IgG-coated ones. We report recruitment of Mannosidase-II positive Golgi-derived vesicles during uptake of diverse targets including latex beads, *E. coli*, *Salmonella* Typhimurium and *Mycobacterium tuberculosis* in human and mouse macrophages. The recruitment of Mannosidase-II vesicles was an early event mediated by focal exocytosis and coincided with the recruitment of transferrin receptor, VAMP3 and dynamin-2. Brefeldin A treatment inhibited Mannosidase-II recruitment and phagocytic uptake of serum coated or uncoated latex beads and *E. coli*. However consistent with previous studies, Brefeldin A treatment did not affect uptake of IgG-coated latex beads. Mechanistically recruitment of Mannosidase-II vesicles during phagocytic uptake required Ca^2+^ from both extra and intra-cellular sources apart from PI3Kinase, microtubules and dynamin-2. Extracellular Ca^2+^ via voltage-gated Ca^2+^ channels establish a Ca^2+^-dependent local PIP3 gradient, which guides the focal movement of Golgi-derived vesicles to the site of uptake. We confirmed Golgi-derived vesicles recruited during phagocytosis were secretory vesicles as their recruitment was sensitive to depletion of VAMP2 or NCS1 however recruitment of recycling endosome marker VAMP3 was unaffected. Both VAMP2 and NCS1 depletion individually resulted in the reduced uptake by macrophages. Together the study provides a previously unprecedented role of Golgi-derived secretory vesicles in phagocytic uptake, the key innate defense function.

## Introduction

Phagocytosis lies at the core of innate defense mechanisms in higher eukaryotes. The process of phagocytosis involves internalization of external particles including pathogens, cellular debris etc. into a specialized membrane bound organelle called phagosomes. The phagosome thus formed undergoes a series of fusion and fission events leading to acquisition of hydrolytic enzymes and microbicidal properties to mature into phago-lysosomes (1-3).

It was perceived initially that the membrane for phagosomes could be solely derived from the plasma membrane (4). However in macrophages, the professional phagocytes, which can engulf objects bigger than their own size without significantly altering their function, there is no apparent decline in the membrane surface area following phagocytosis(5). Rather capacitance measurement experiments showed that a concomitant increase in the membrane surface area precedes resealing of the phagosome, suggesting supply of phagosome membrane from intracellular sources(5). Professional phagocytes are therefore expected to be under tremendous pressure to sustain the supply of the membrane so as to effectively phagocytize their targets. Consequently a variety of membrane bound organelles including the vesicles originating from recycling endosomes and lysosomes as well as ER were later shown to provide membrane for the nascent phagosome(6-8).

Vesicle recruitment during phagocytosis involves exocytosis of vesicles in the vicinity of nascent phagosomes (9). Studies in the past have shown secretion of vesicular contents like lysosomal hydrolases and azurophil granules during engulfment by macrophages and neutrophils respectively, thereby coupling the process of phagocytosis with targeted exocytosis, a process also termed as focal exocytosis (7,9). Recycling endosome marker VAMP3, a soluble Nethylmaleimide-sensitive factor attachment protein receptor proteins on vesicles (v SNARE), was shown to accumulate at the early phagosomes through focal exocytosis (7). Moreover Synaptotagmin V (SytV), a Ca^2+^ sensor on the recycling endosomes was shown to regulate phagocytosis but not phagosome maturation, suggesting a key role of Ca^2+^ in the regulation of vesicle exocytosis (10). Similarly SytVII, a lysosome resident Ca^2+^ sensor was reported to regulate delivery of lysosome membrane to the nascent phagosomes (11). Interestingly, SytVII was also shown to regulate Ca^2+^ dependent exocytosis of lysosomes during plasma membrane repair (12). Recently TRPML1, a key Ca^2+^ channel in the lysosomes was shown to regulate focal exocytosis during phagocytosis of large particles (13). Phosphoinositide3kinase (PI3K) is yet another key regulator of phagocytosis, and is believed to help pseudopod extension around the particles during phagocytosis (14). However, inhibition of PI3K by wortmannin may not limit membrane availability for phagocytosis (14). There are some reports however suggesting the role of PI3K in recruiting the membranes from intracellular sources (15). Yet another study revealed recruitment of endoplasmic reticulum (ER) at the site of phagocytosis to provide membrane for the newly formed phagosomes, a process that was regulated by the ER resident SNARE protein ERS24 (2,6).

Intriguingly, despite being intricately involved with the recycling endocytic network and as one of the major sources of vesicles destined to the plasma membrane like secretory vesicles, involvement of Golgi Apparatus (GA) or vesicles derived from GA in the process of phagocytosis has been ruled out (16). However in an interesting observation, fragmentation and reorganization of GA was reported during frustrated phagocytosis (17). Interestingly majority of the reports that conclude no direct involvement of GA during phagocytic uptake relied entirely on the experiments that were limited to Fcγ-Receptor mediated phagocytosis (18-20). It is important to note that macrophages employ several independent receptor systems for phagocytosis including complement mediated, pattern receptor mediated and scavenger receptor mediated phagocytosis (3,21). Whether Golgi apparatus is dispensable even for these phagocytic pathways remains unknown.

Here we studied the role of vesicles derived from the GA during phagocytic uptake using Mannosidase-II as the marker for these vesicles. Mannosidase-II, a medial and trans-Golgi marker was also detected within secretory granules and at the surface of certain cell types like enterocytes, pancreatic acinar cells and goblet cells (22). Later, using Mannosidase-II as an exclusive marker of Golgi-derived vesicles its recruitment to cell surface was used to track the role of these vesicles during plasma membrane repair in mouse bone marrow derived primary macrophages (22,23). Here we show, Mannosidase-II containing Golgi-derived vesicles are recruited during phagocytosis of a variety of targets including inert particles (latex beads, uncoated or serum coated), non-pathogenic bacteria (*E. coli*) and pathogenic bacteria (*Salmonella* Typhimurium and *Mycobacterium tuberculosis*) in mouse (RAW264.7 and BMDMs) and human (THP-1 and U937) macrophages. The recruitment of Mannosidase-II vesicles at the site of phagocytosis occurred through the process of focal exocytosis. This process was dependent on voltage-gated Ca^2+^ channels that helped establish a Ca^2+^ dependent PIP3 gradient for uptake. Mechanistically, TGN and secretory vesicle resident protein NCS1 sensed and triggered the movement of vesicles to help phagosome biogenesis, which was also dependent on secretory vesicle specific v-SNARE VAMP2. We trace here hitherto unknown function of Golgi-derived vesicles during phagocytic uptake in macrophages and also provide mechanistic basis for the same.

## Results

### *α*-Mannosidase-II (MAN2A) localizes to Golgi Apparatus (GA) and Golgi derived vesicles

To confirm the localization of Mannosidase-II to GA in our experimental set-up, we nucleofected THP-1 human macrophages and RAW264.7 mouse macrophages with a mCherry-tagged Mannosidase-II construct [mCherry-MannII-N-10 (mCherry-MAN2A), Addgene plasmid # 55074] and visualized the cells under Nikon A1R confocal microscope (see methods). Expression of Mannosidase-II in these cells distinctly labeled the GA as shown in figure 1A. We also immuno-stained Mannosidase-II using specific antibody, which labeled the GA (Fig. 1B). Treatment of mCherry-MAN2A expressing cells with Brefeldin A, the classical inhibitor of Golgi function, resulted in the disruption and fragmentation of the GA structure (Fig 1C). In both transfected as well as antibody stained cells, in addition to the main GA we could also observe several smaller punctate structures, representing the Golgi-derived vesicles (Fig. 1A and B). In U937 cells stably expressing mRuby-tagged Beta-1,4 Galactosyltransferase (B4GALT1-mRuby), a mid-Golgi resident enzyme, such vesicle like structures could be rarely seen (Fig. 1D), Mannosidase-II however co-localized with B4GALT1-mRuby (Fig. 1D). These results confirmed that the Mannosidase-II construct used in this study labeled both the GA and Golgi-derived vesicles.

**Figure 1:**
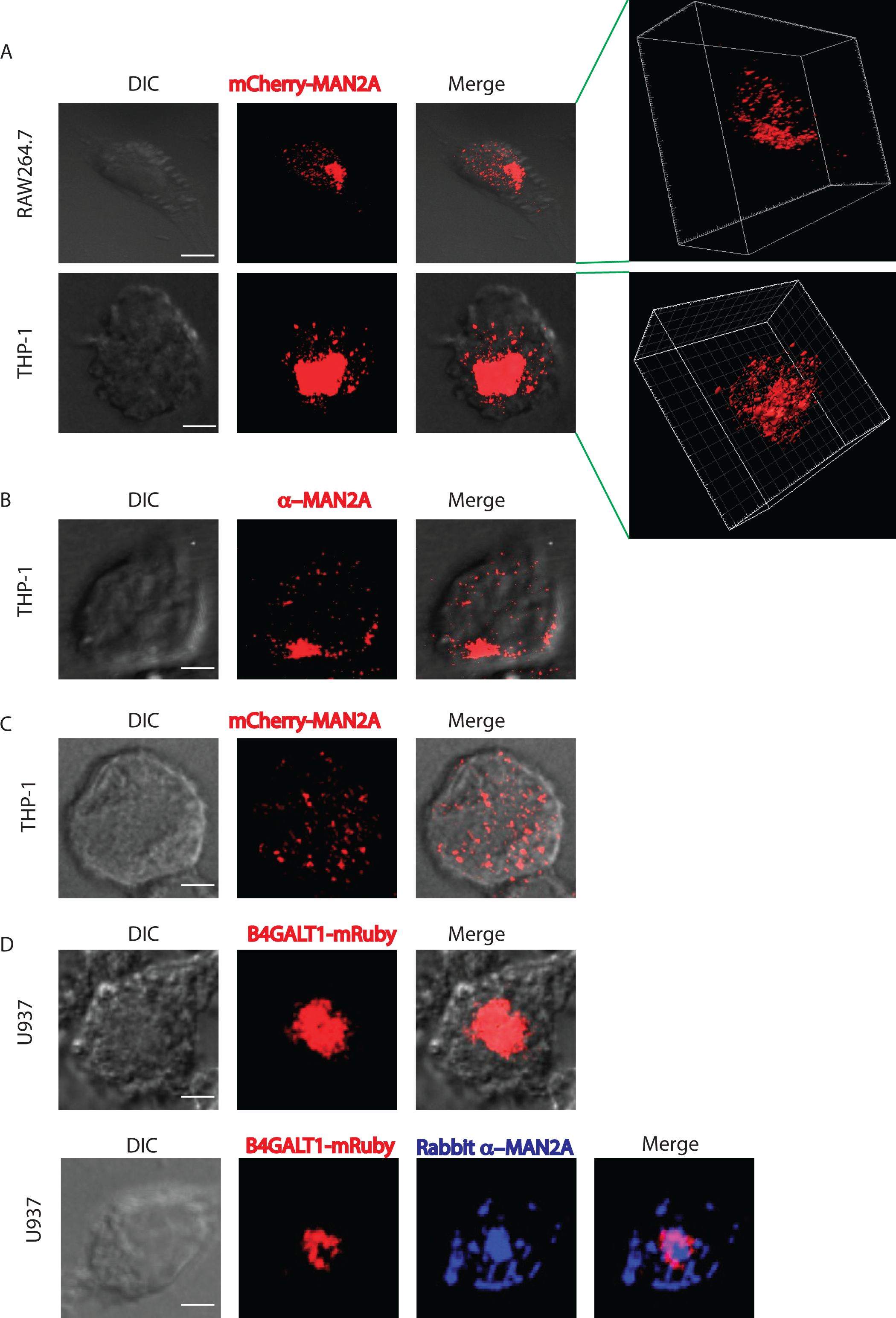
Mannosidase-II is a marker of Golgi apparatus and vesicles derived from the Golgi. A) RAW264.7 cells and THP-1 derived macrophages were nucleofected with mCherry-MAN2A. At 24 hours post nucleofection, cells were visualized under the confocal microscope (see methods for detail). The 3-D plots at the right were created using Imaris 7.2 software tool (scale bar: 5μm). B) THP-1 derived macrophages were permeabilized using 0.5% TritonX-100 and stained with anti-Mannosidase-II antibody followed by the secondary antibody tagged with Alexa-560, fixed and visualized under the confocal microscope (scale bar: 5μm). C) THP-1 derived macrophages were nucleofected with mCherry-MAN2A. At 24 hours post nucleofection, the cells were treated with Brefeldin A (20μM) for 4 hours and visualized under the microscope (scale bar: 5μm). D) U937 human macrophages, stably expressing the Golgi marker mRuby-B4GALT1 (red, β-galactosyl transferase). For the lower panel, B4GALT1 (red) expressing U937 macrophages were stained with anti-Mannosidase-II antibody followed by secondary antibody (Alexa-405, blue, scale bar: 5μm).

### Mannosidase-II is recruited at the site of phagocytic uptake

RAW264.7 cells transfected with mCherry tagged Mannosidase-II were incubated with mouse serum coated latex beads (see methods). At 30 minutes and one-hour post addition of beads to the macrophages, cells were fixed and visualized under the microscope. We observed the recruitment of Mannosidase-II to the sites where latex beads were phagocytosed, forming ring like structure around the beads at many instances (Fig. 2A). Overall intensity of Mannosidase-II at the bead surface had a median of ~900 at 30 minutes and ~1500 arbitrary units (A.U.) at 1 hour as determined using the 3-D module tool in Imaris 7.2 (see methods, Fig 2A). To check whether Mannosidase-II recruitment during phagocytosis was a general phenomenon we also monitored uptake of non-pathogenic bacterium *E. coli* and pathogenic bacteria *Salmonella* Typhimurium. In both the cases, we could see Mannosidase-II recruited at very early stages of phagocytosis. In the case of *E. coli,* nearly 60% of the bacteria at the site of entry showed Mannosidase-II recruitment at 15 and 30 minutes post-infection while nearly 30% of *Salmonella* containing phagosomes showed Mannosidase-II recruitment at 5 and 10 minutes (Fig 2B). In the case of yet another pathogenic bacteria *Mycobacterium tuberculosis* strain H37Rv, Mannosidase-II was recruited at 30-40% phagosomes at the time of entry (Fig. 2C). Recruitment of Mannosidase-II at the early phagosomes in case of H37Rv was also verified by immuno-staining against Mannosidase-II (Fig. 2D). We also verified the recruitment of Mannosidase-II in mouse bone marrow derived macrophages (BMDMs) during phagocytic uptake of *E. coli*, thus ensuring that it was not a phenomenon restricted only to the cell-lines (Fig. 2E). To confirm that the Mannosidase-II positive phagosomes indeed represented early stages of phagosomes, we immuno-stained the mCherry-MAN2A expressing cells, that were either incubated with mouse serum coated latex beads or infected with *Mtb*, with anti-transferrin receptor (TfR) antibody. We observed significant overlap between recruited Mannosidase-II and TfR around the internalized beads and *Mtb* (Fig. 3A and 3B). TfR recruitment around phagocytized beads showed saturating levels with median of around 4095 A.U., much higher than the intensity distribution of Mannosidase-II (~1000 A.U., Fig. 2D). In case of *Mtb* too, TfR showed relatively higher co-localization with the phagosomes (~40-60%) compared to Mannosidase-II (30-50%, Fig. 3B). Expectedly, TfR on *Mtb* phagosomes declined rapidly from one-hour post infection to two hours post infection (Fig. 3B).

**Figure 2:**
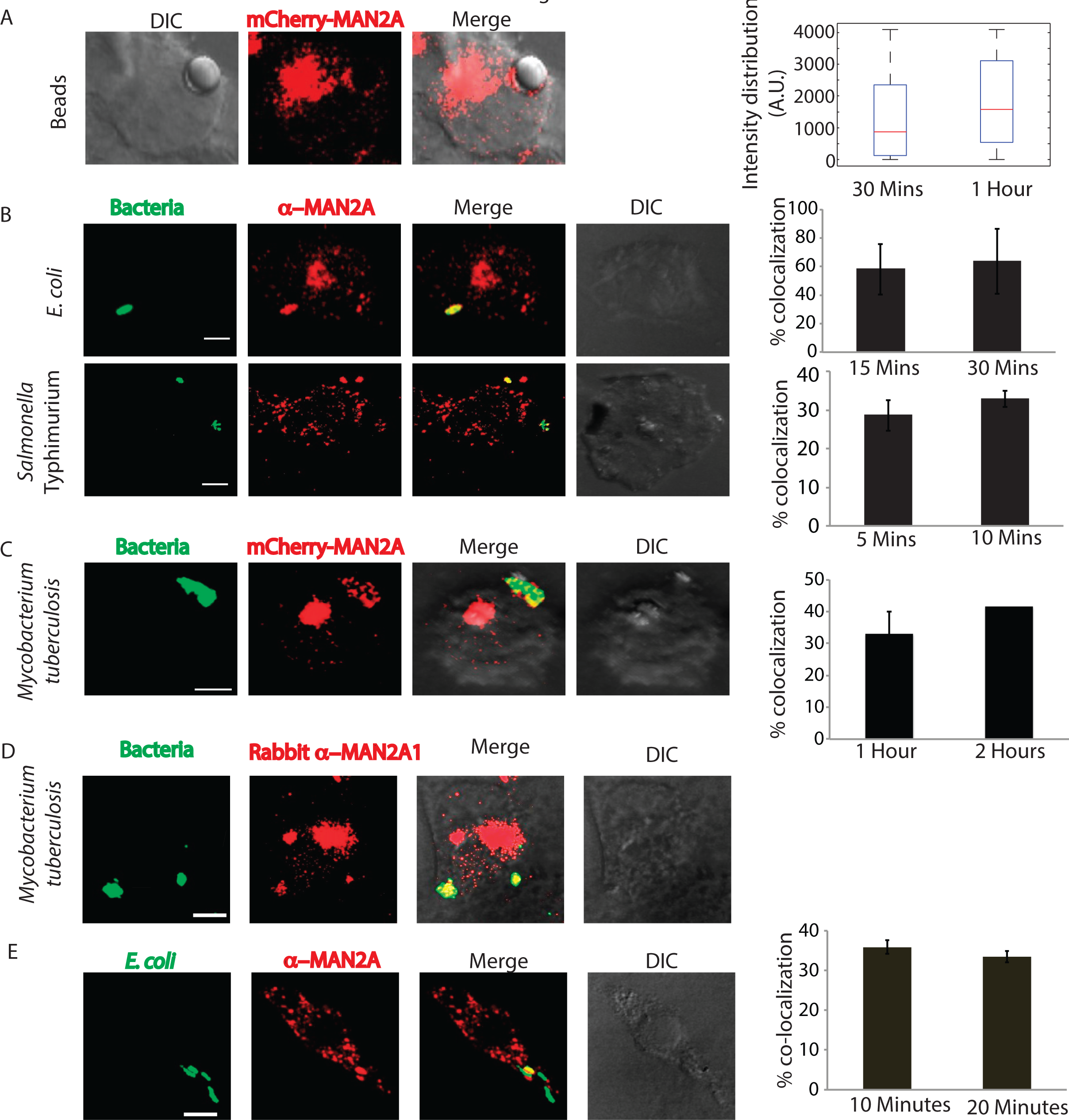
Mannosidase-II is recruited at the site of phagocytosis. A) RAW264.7 macrophages expressing mCherry-MAN2A (red) were incubated with mouse serum coated latex beads for 30 minutes and 1 hour. The images shown are representative from the 30 minutes samples. At the right, the total intensity of mCherry-MAN2A puncta on the bead surface was estimated using 3D spot creation module in Imaris 7.2 and the intensity distribution of the population has been plotted (see methods for detail). The box plot represents data from more than 200 beads analyzed from two different experiments (values ± S.D.; scale bar: 4μm). B) THP-1 derived macrophages were infected with GFP expressing *E. coli* (green, top panel) or *Salmonella* Typhimurium (green, lower panel) for 15 and 30 minutes. Cells were then stained with anti-Mannosidase-II antibody followed by Alexa 568 tagged secondary antibody (red). Images shown are representative from the 15 minutes samples from both the experiments. The bar plots at the right show % co-localization of *E. coli* or *Salmonella* with Mannosidase-II at the surface at 5 and 10 or 15 and 30 minutes respectively post infection. Data represents average of more than 200 bacteria from two different experiments (values ± S.D.; scale bar: 3μm). C) mCherry-MAN2A (red) expressing THP-1 derived macrophages were infected with PKH67 labeled H37Rv (green) for 1 and 2 hours. Samples were fixed and visualized under the microscope. The bar plot at the right shows % co-localization of H37Rv with Mannosidase-II at both these time points. Data represents average of more than 200 bacteria from two different experiments (values ± S.D.; scale bar: 4μm). D) THP-1 derived macrophages were infected with PKH67 labeled H37RV and fixed at respective time points. These cells were then permeabilized using 0.05% TritonX-100 and stained with anti Mannosidase-II antibody followed by the secondary antibody tagged with Alexa-568, fixed and visualized under the confocal microscope (scale bar: 3μm). E) Mouse BMDMs were infected with GFP expressing *E coli* for 10 minutes, fixed and stained with anti-MAN2A antibody (scale bar: 3μm).

### Characterization of early phagosomes reveals direct involvement of secretory vesicles derived from the GA in phagosome biogenesis

As shown above and by others (22,23), Mannosidase-II marks both GA and Golgi-derived vesicles. In most of the fields observed in the previous sections, we could see Mannosidase-II organized into a separate Golgi structure, underscoring the fact that the phagosome associated Mannosidase-II were derived from the recruitment of Golgi-derived vesicles at the site of infection. To further confirm the selective enrichment of Mannosidase-II at the nascent phagosomes, we purified latex beads phagosomes using sucrose density gradient ultra-centrifugation from macrophages within 1 hour of uptake (Fig. 3C). The phagosome preparation showed presence of expected early phagosome markers like TfR and RAB5 (Fig. 3D). They also showed presence of previously reported markers like VAMP3 from recycling endosomes and Calnexin from ER (Fig. 3D). At the same time, the phagosome preparation was devoid of any mitochondrial, nuclear or cytosolic contamination (Fig. 3D). In agreement with the microscopy data, phagosomes also showed presence of Mannosidase-II, the marker for Golgi-derived vesicles (Fig. 3D). In addition, we could also score the presence of another Golgi-derived secretory vesicles marker NCS1 in the phagosomes (Fig. 3D). We verified that there was no GA contamination in the preparation as the phagosomes were devoid of B4GALT1 (Fig. 3D).

**Figure 3:**
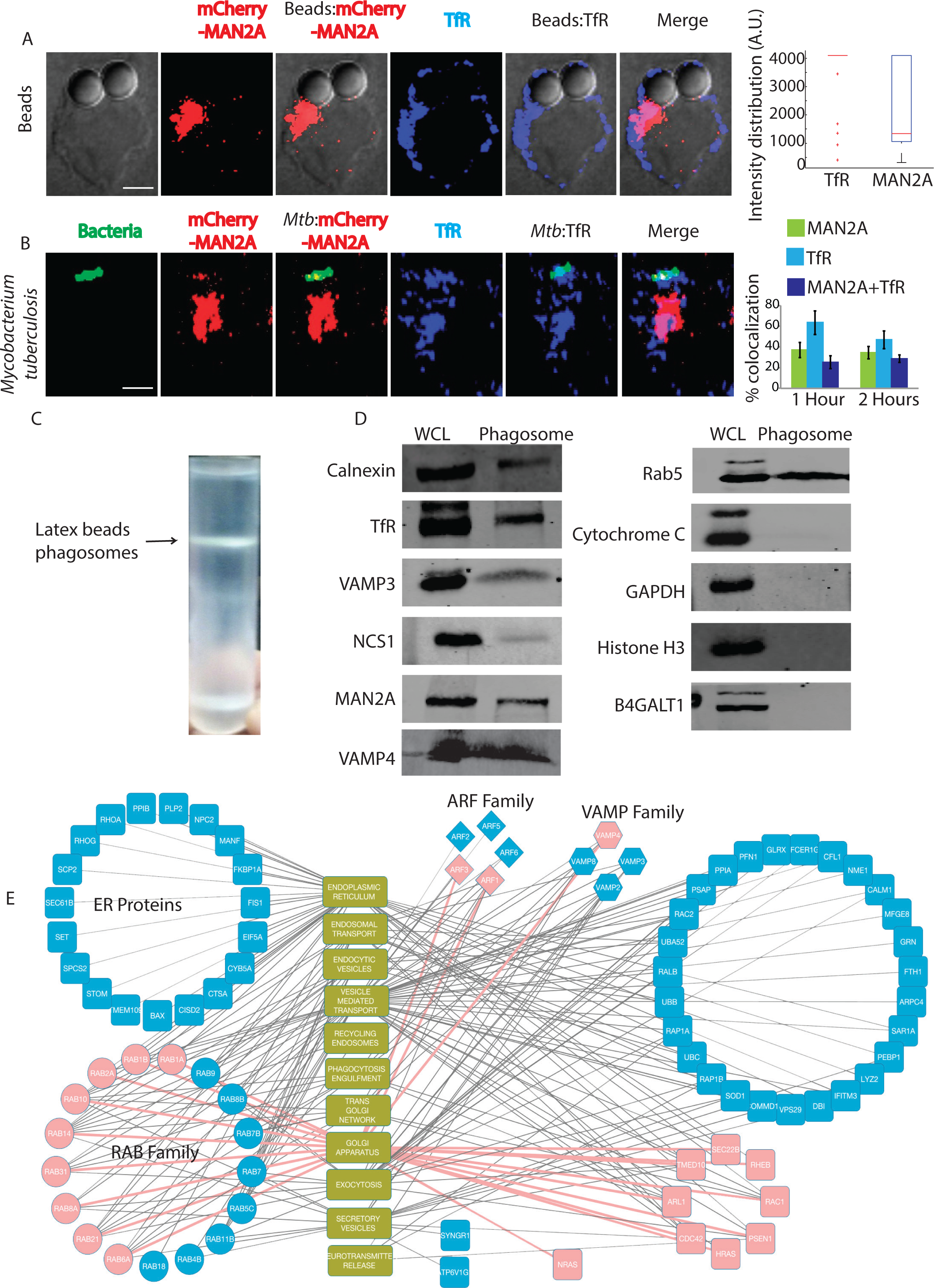
Early recruitment of Mannosidase-II at phagosomes and proteomic characterization. A) mCherry-MAN2A (red) expressing RAW264.7 macrophages were incubated with mouse serum coated latex beads for 30 minutes or 1 hour. At the respective time points, cells were immune-stained with anti-TfR antibody followed by a secondary antibody tagged with Alexa 405 (blue). Presence of TfR or Mannosidase-II at the bead surface was calculated in terms of fluorescence intensity using the 3D spot creation module in Imaris 7.2software. The box-plot at the right shows data from more than 100 beads from two independent experiments (scale bar: 5μm). B) mCherry-MAN2A (red) expressing RAW264.7 macrophages were infected with PKH67 labeled H37Rv (green) for 1 and 2 hours. At the respective time points, samples were fixed and stained with anti-Transferrin receptor antibody followed by Alexa-405 tagged secondary antibody (blue). The images are representative from the 1 hour time point. For the plots at the right, % co-localization of H37Rv with Mannosidase-II, TfR or both Mannosidase-II and TfR was calculated using Imaris 7.2. The data represents average of more than 150 bacteria from three different experiments (values ± S.D.; scale bar: 5μm). C) Preparation of latex bead phagosomes from THP-1 derived macrophages on a sucrose density gradient (see methods) D) THP-1 derived macrophages were incubated with latex beads (1μm size) for 1 hour. Phagosomes were isolated using differential density ultracentrifugation and samples were probed for indicated markers using Western blots. E) Latex beads phagosomes isolated from RAW264.7 macrophages were lysed and resolved on a 10% SDS-PAGE. The lane below 25kDa molecular weight was analyzed using mass spectrometry to identify enrichment of low molecular weight proteins (see methods). The list of genes identified was then searched in the AMIGO2.0 database to establish functional association. Finally the representative network was constructed using Cytoscape 2.6.1. The pink nodes and edges in the network denote association with the Golgi apparatus.

Trafficking of vesicles in the cells is regulated by a large number of proteins including small GTPases like RABs, ADP-Ribosylation factors (ARFs) and SNAREs (25-27). In order to understand which RABs, SNAREs or ARFs could be involved in the recruitment of Mannosidase-II vesicles at the phagosomes we performed mass-spectrometry of the purified latex beads phagosome preparations, specifically to identify molecules below 25kDa molecular weight (see methods). We were able to identify about 290 proteins from this preparation, all of them below 25kDa molecular weight (Supplementary table S1). We performed a gene ontology analysis on these proteins using Amigo2.0 database to specifically see enrichment of eleven classes including ER, GA, secretory vesicles, recycling endosomes, endocytic vesicles, neurotransmitter release, TGN, vesicle mediated transport, phagocytosis/engulfment, exocytosis and endosomal transport (Fig. 3E, Supplementary table S2). A large number of proteins belonging to ER were found in the phagosomes (Fig. 3E). The phagosome mass spectrometry data revealed presence of 17 different RABs, 5 out of 6 known ARFs and four VAMPs. Many of these proteins are involved with Golgi apparatus, exocytosis, secretory vesicles and neurotransmitter release (Fig. 3E). VAMP2 gets enriched in the secretory vesicles that are released from the GA and targeted to the membranes (28). It also helps in the release of neurotransmitters by regulating the exocytosis of secretory vesicles (29). Collectively presence of markers associated with vesicles derived from the GA reconfirmed the utility of this compartment as an additional source of membrane for phagosome biogenesis.

### Requirement of Golgi apparatus during phagocytosis

Having observed overwhelming representation of proteins from GA in the phagosome proteome and clearly established the recruitment of MAN2A positive Golgi-derived vesicles at the phagosomes, it was important to test whether GA could be directly involved in regulating the uptake. It must be noted that classically role of GA during Fcγ-receptor mediated phagocytosis has been ruled out (16). Corroborating previous studies, we observed that uptake of human-IgG coated latex beads was unaffected by Brefeldin A treatment, an established Golgi inhibitor at doses ranging from 10, 20 30 to 100μg/ml (Fig. 4A). However uptake of serum-opsonized latex beads, latex beads opsonized with complement deactivated serum or uncoated latex beads got significantly inhibited by Brefeldin A treatment at doses as low as 10 and 20 μg/ml respectively (Fig. 4B, C and D). Similarly uptake of *E. coli* was also significantly inhibited by Brefeldin A treatment at 30 and 100μg/ml (Fig. 4E). Interestingly, Brefeldin A treatment also led to diminished recruitment of Mannosidase-II at the latex bead phagosomes or *E. coli* phagosomes (Fig. 4F and G). Mannosidase-II recruitment was not entirely absent in case of Fcγ-receptor mediated phagocytosis, was sensitive to Brefeldin A treatment but, unlike other cargos, that did not lead to any affect on uptake (Fig. 4H). Since requirement of Golgi function varied between different modes of phagocytosis, we questioned that should not reflect the effect of Brefeldin A on the corresponding receptor expression level at the cell surface. All cell surface receptors travel through the Golgi and their expression levels may very much get perturbed in Brefeldin A treated cells. However none of the receptors tested including Fcγ-receptor, complement receptor (CR3), mannose receptor and Dectin-2 showed any noticeable decline at the doses and duration of treatment used for the uptake studies (Fig. 4I).

**Figure 4:**
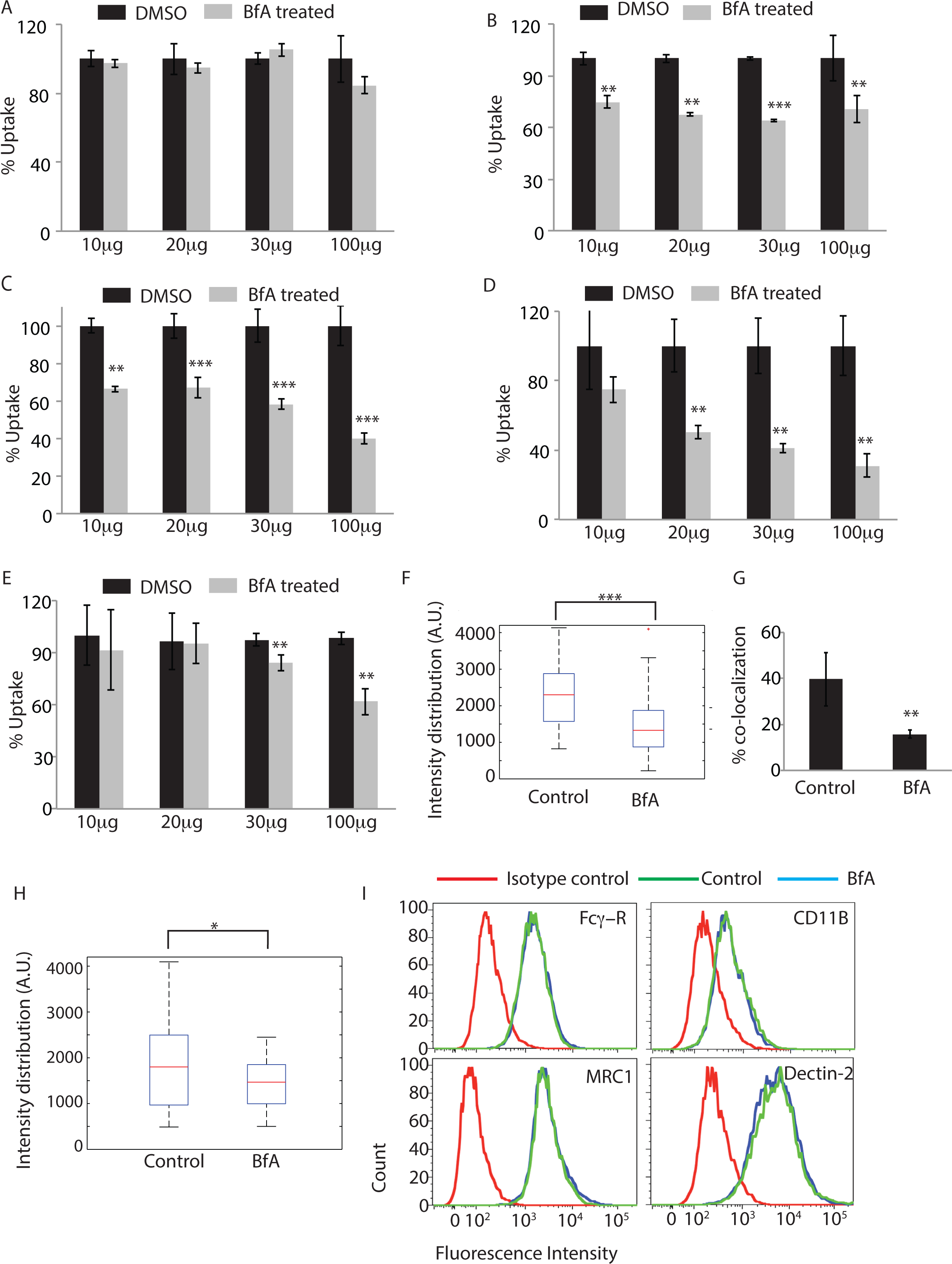
Role of Golgi apparatus during phagocytosis. THP-1 macrophages were incubated with human IgG-coated FITC-labeled latex beads (A), serum-coated FITC-labeled latex beads (B), complement deactivated serum coated latex beads (C), uncoated latex beads (D) or *E. coli* (E) for 30 minutes in the absence or presence of Brefeldin A at concentrations mentioned. BfA treatment was given 30 minutes prior to the addition of the beads or *E. coli*. Samples were acquired in a flow cytometer to measure uptake. Plots represent data from average of three independent experiments. Error bars represent SD (**p‐ value<0.01; ***p-value<0.005). (F) THP-1 macrophages were allowed to uptake serum-coated FITC-labeled latex beads for 30 minutes in the presence or absence of BfA. Samples were fixed and stained for Mannosidase-II using anti-MAN2A antibody followed by alexa-405 secondary antibody. Confocal images acquired were analyzed using 3-D module in Imaris 7.2 software to calculate Mannosidase-II intensity distribution over the latex beads (***p-value<0.001). (G) GFP expressing *E. coli* were incubated with THP-1 macrophages for 30 minutes in the presence or absence of BfA. Samples were fixed and stained for Mannosidase-II using anti MAN2A antibody followed by alexa-405 secondary antibody. Confocal images were analyzed using Imaris software to calculate percent co-localization. (H) THP-1 macrophages were allowed to uptake IgG-coated FITC-labeled latex beads for 30 minutes in the presence or absence of BfA. Samples were fixed and stained for Mannosidase-II using anti-MAN2A antibody followed by alexa-405 secondary antibody. Confocal images acquired were analyzed using 3-D module in Imaris software to calculate Mannosidase-II intensity distribution over the latex beads. co-localization (*p-value<0.05). I) U937 derived macrophages were incubated with 20ug/ml of Brefeldin A for 60 mins, fixed and stained for receptors mentioned. Cells were acquired by flow cytometry and overlays analyzed using Flow Jo.

### Golgi-derived vesicles co-operate with the vesicles derived from the recycling endosomes during phagocytosis

Recruitment of vesicles from endocytic origin (VAMP3 positive) and lysosomal origin (LAMP1 positive) during phagocytosis has been reported earlier (7,11). We also observed presence of VAMP3 on the early latex beads phagosome proteome as shown above (Fig. 3D). We verified that VAMP3compartment represented distinct population of vesicles than those represented by Mannosidase-II (Fig. 5A). However, since VAMP3 is known to traffic through the Golgi apparatus, there were some overlaps between Mannosidase-II and VAMP3 within the Golgi stacks (Fig. 5A). However outside the GA, there was very limited overlap between Mannosidase-II and VAMP3. In mCherry-MAN2A expressing RAW264.7 cells, both latex beads and *Mtb* very early during phagocytosis (30 minutes and 1 hour for beads; 1 hour and 2 hours for *Mtb*) showed very high recruitment of and co-localization with VAMP-3 (Fig. 5B and C). In case of *Mtb*, more than 90% of the Mannosidase-II positive phagosomes were also positive for VAMP-3 (Fig. 5C). Thus spatially and temporally, Mannosidase-II recruitment at the phagosomes was strikingly similar to VAMP3 recruitment. Interestingly, unlike Mannosidase-II recruitment, VAMP3 recruitment was not sensitive to Brefeldin A treatment during uptake of latex beads or *E. coli* (Fig. 5D and E respectively). The selective influence of Brefeldin A on Mannosidase-II recruitment suggests independent pathways for the recruitment of Mannosidase-II vesicles and VAMP3 vesicles. While Mannosidase-II vesicles are known to originate from Golgi apparatus, VAMP3 vesicles are recruited from recycling endosomes (7). Thus it seems vesicles from different origins co-operate in aiding the biogenesis of phagosomes.

**Figure 5:**
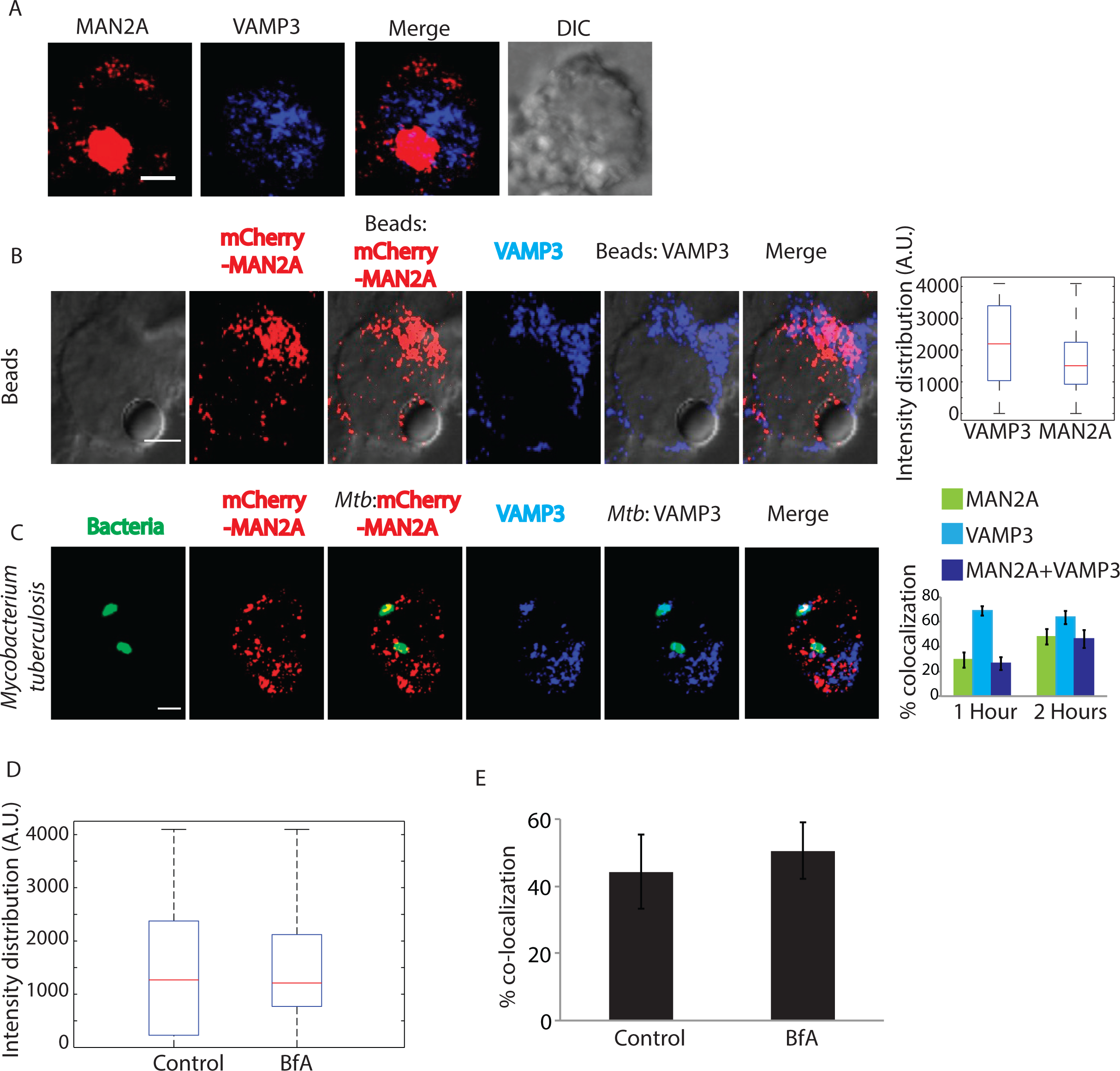
Vesicles derived from Golgi apparatus are distinct from recycling endosome vesicles and are recruited independently. (A) U937 cells expressing MAN2A mCherry were fixed and stained with anti VAMP3 antibody followed by alexa-405 labeled secondary antibody (Scale bar-4μm). (B) mCherry-MAN2A (red) expressing RAW264.7 macrophages were incubated with mouse serum coated latex beads for 30 minutes or 1 hour. At the respective time points, cells were immune-stained with anti VAMP-3 antibody followed by a secondary antibody tagged with Alexa 405 (blue). Presence of VAMP-3 or Mannosidase-II at the bead surface was calculated using the 3D spot creation module in Imaris 7.2 software. The box-plot at the right shows data from more than 100 beads from two independent experiments (scale bar: 5μm). (C) mCherry-MAN2A (red) expressing THP-1 derived macrophages were infected with PKH67 labeled H37Rv (green) for 1 and 2 hours. At the respective time points, samples were fixed and stained with anti-VAMP-3 antibody followed by Alexa-405 tagged secondary antibody (blue). The images are representative from the 1hour time point. For the plots at the right, % co-localization of H37Rv with Mannosidase-II, VAMP-3 or both Mannosidase-II and VAMP-3 was calculated using Imaris 7.2. The data represents average of more than 150 bacteria from three different experiments (values ± S.D.; scale bar: 4μm). (D) U937 cells were incubated with serum-coated FITC-labeled latex beads in the presence or absence of BfA for 30 minutes. Samples were fixed and stained with anti-VAMP3 antibody followed by alexa-405 labeled secondary antibody. Images were analyzed using 3-D module in Imaris 7.2 and VAMP3 intensity distribution in the latex beads were calculated. (E) U937 cells were incubated with GFP expressing *E. coli* in the presence or absence of BfA for 30 minutes. Samples were fixed and stained with anti-VAMP3 antibody followed by alexa-405 labeled secondary antibody. Images were analyzed using 3-D module in Imaris 7.2 and VAMP3 intensity distribution in the latex beads were calculated.

### Mannosidase-II positive Golgi-derived vesicles are recruited at the phagosomes through focal exocytosis

We next wanted to understand how Mannosidase-II vesicle recruitment was regulated during phagocytosis. Previous studies show the recruitment of VAMP-3 positive vesicles from the recycling endosomes is dependent on focal exocytosis (7). It was previously reported that Mannosidase-II positive Golgi-derived vesicles get recruited at the site of membrane damage during repair (23). It is also well established that vesicular movement during membrane repair occurs through focal exocytosis (12). One of the key regulators of focal exocytosis, specifically for post-Golgi transport vesicles towards the plasma membrane, is dynamin (30). Dynamins are the member of large GTPase family and dynamin-2 is the ubiquitously expressed isoform while dynamin-1 is mostly neuronal (31,32). Their involvement in budding of transport vesicles from the Golgi is also well documented along with their role in regulating focal exocytosis (13,33). Additionally dynamins are also involved in the endocytosis where they regulate scission of endocytic vesicles from the membrane (Liu, 2011, Traffic, 12, 1620). In mCherry-MAN2A expressing RAW264.7 cells, we observed significant overlap of Mannosidase-II with dynamin-2 on the phagosomes containing either latex beads or *Mtb* (Fig. 6A and 6B). Again as in the case of TfR, dynamin-2 recruitment on the nascent phagosomes was relatively higher than Mannosidase-II recruitment for both latex beads and *Mtb* (Fig. 6A and 6B). Presence of dynamin-2 at the phagosomes was consistent with previous reports, which showed their recruitment at early phagosomes during phagocytosis (34).

Dynamin mediated focal exocytosis can be inhibited by dynasore, a specific dynamin inhibitor (13). We next inhibited dynamin-2 by dynasore treatment for varying period of time and observed its effect on Mannosidase-II recruitment and phagocytosis. In the case of latex beads, four hours of pre-treatment with dynasore (at 40 and 80μM) resulted in a significant inhibition in the recruitment of Mannosidase-II at the phagosomes (Fig. 6C). It also markedly reduced the phagocytosis of latex beads (Fig. 6D). Dynasore treatment also inhibited uptake of *Mtb* in THP-1 macrophages (Fig. 6E). There was noticeable decline in the recruitment of Mannosidase-II at the *Mtb* phagosomes (Fig. 6F). The reduction in uptake upon dynasore treatment was not due to the inhibition of endocytic functions of dynamins since dynasore unlike Mannosidase-II, did not inhibit recruitment of recycling endosome marker VAMP3 to the phagosomes. Dynamin assisted movement of vesicles also require microtubules (30). Inhibition of microtubules by nocodazole treatment resulted in a transient decline in the uptake of *Mtb* and latex beads (Fig. 6G and 6H). Further, nocodazole treatment resulted in a marked decline in the recruitment of Mannosidase-II to the phagosomes containing *Mtb* and latex beads (Fig. 6I and 6J). These results together established that the Mannosidase-II positive Golgi-derived vesicles were recruited at the phagosomes through focal exocytosis and were assisted by dynamins.

**Figure 6:**
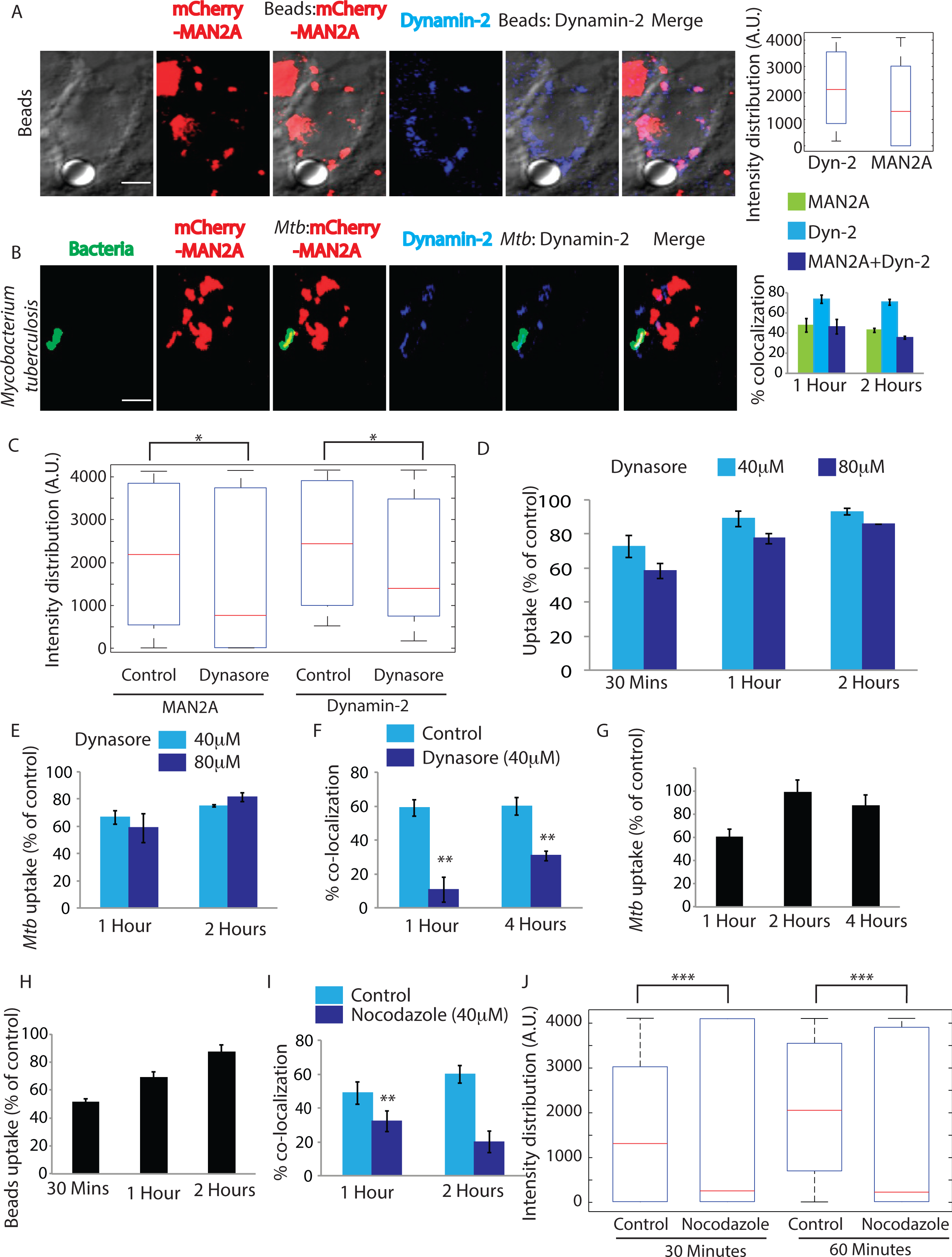
Golgi-derived vesicles are recruited through focal exocytosis. A) mCherry-MAN2A (red) expressing RAW264.7 macrophages were incubated with mouse serum coated latex beads for 30 minutes or 1 hour. At the respective time points, cells were immune-stained with anti-dynamin2 antibody followed by a secondary antibody tagged with Alexa 405 (blue). Presence of dynamin2 (Dyn-2) or Mannosidase-II at the bead surface was calculated using the Imaris 7.2 software. The box-plot at the right shows data from more than 100 beads from two independent experiments (values ± S.D.; scale bar: 4μm). **B**) mCherry-MAN2A (red) expressing THP-1 macrophages were infected with PKH67 labeled H37Rv (green) for 1 and 2 hours. At the respective time points, samples were fixed and stained with anti-dynamin2 antibody followed by Alexa-405 tagged secondary antibody (blue). The images are representative from the 1hour time point. For the plots at the right, % co-localization of H37Rv with Mannosidase-II, dynamin2 or both Mannosidase-II and dynamin2 was calculated using Imaris 7.2. The data represents average of more than 150 bacteria from three different experiments (values ± S.D.; scale bar: 4μm). C) mCherry-MAN2A (red) expressing RAW264.7 macrophages were pretreated with 80μM Dynasore and incubated with mouse serum coated latex beads for 30 minutes or 1 hour. At 30 minutes, samples were fixed and stained with anti-dynamin2 antibody followed by Alexa-405 tagged secondary antibody (blue). Intensity of Mannosidase-II and dynamin at the bead surface was calculated using the 3D spot creation module in Imaris 7.2 software. The box-plot shows data from more than 100 beads from two independent experiments (*p-value<0.05). D) THP 1 derived macrophages were treated with the respective dynasore concentrations for 4 hours and incubated with FITC labeled latex beads (1μm). At the respective time points cells were fixed and analyzed by flow cytometry. To calculate % uptake, data for each cargo for a given time point, the dynasore treated set was normalized against the respective untreated control set (values ± S.D.). E) THP-1 derived macrophages were pre-treated with 40μM or 80μM Dynasore for 4 hours and infected with PKH67 labeled H37RV. The cells were fixed at respective points and analyzed by flow cytometry (values ± S.D.). F) THP-1 derived macrophages expressing mCherry-MAN2A were pretreated with 40μM Dynasore for 4 hours and infected with PKH67 labeled H37RV. The cells were fixed at respective time points and analyzed by confocal microscope. The percent co-localization was determined by Imaris 7.2. Data represents average of more than 200 bacteria from two different experiments (values ± S.D.,**p‐ value<0.01)). G) THP1 derived macrophages were pre-treated with 25μM Nocodazole for 2.5 hours and infected with PKH67 labeled H37RV. The cells were fixed at respective points and analyzed by flow cytometry (values ± S.D.). H) THP1 derived macrophages expressing mCherry-MAN2A were pretreated with 25μM Nocodazole for 2.5 hours and infected with 1μm-FITC-labelled latex beads. The cells were fixed at respective points and analyzed by flow cytometry (values ± S.D.). I) THP1 derived macrophages expressing mCherry-MAN2A were pretreated with 25μM nocodazole for 2.5hours and infected with PKH67 labeled H37RV. The cells were fixed at respective points and analyzed by confocal microscope. The percent co-localization was determined using Imaris 7.2. Data represents average of more than 200 bacteria from two different experiments (values ± S.D.; **p‐ value<0.01)). RAW264.7 macrophages expressing mCherry-MAN2A were pre-treated with 25μM nocodazole for 2.5 hours and infected with 4μm latex beads. The cells were fixed at respective points and analyzed by confocal microscope. 3D-creation of bead was done to determine the maximum fluorescence intensity of Mannosidase II on the bead, via spot module in Imaris 7.2. The box-plot at the right shows data from more than 100 beads from two independent experiments (**p-value<0.01).

### Phosohatidylinositol3Kinase (PI3K) is required for Mannosidase-II recruitment at the nascent phagosomes and phagocytosis

The role of PI3K in regulating phagocytosis is well known where its function is believed to be mostly required for pseudopod extension around the cargo during phagocytosis (14). We wanted to test whether some of the effects of PI3K on phagocytosis were due to its involvement in the recruitment of Mannosidase-II vesicles for phagocytosis. Inhibition of PI3K by wortmannin led to nearly 80% decline in the uptake of latex beads at 30 and 60 minutes (Fig. 7A). In case of *Mtb*, the decline in uptake was ~85% at 1 and 2 hours post infection and 70% at 4 hours post-infection (Fig. 7B). Curiously, PI3K inhibition also resulted in a marked decrease in the recruitment of Mannosidase-II at the nascent phagosomes (Fig. 7C and 7D). Thus at least to some extent, it seems the effect of inhibition of PI3K on phagocytosis may be due to a reduced recruitment of Mannosidase-II positive Golgi-derived vesicles at the phagosomes. Similar effect of PI3K inhibition on the membrane recruitment for the newly forming phagosomes has been discussed previously (15).

**Figure 7:**
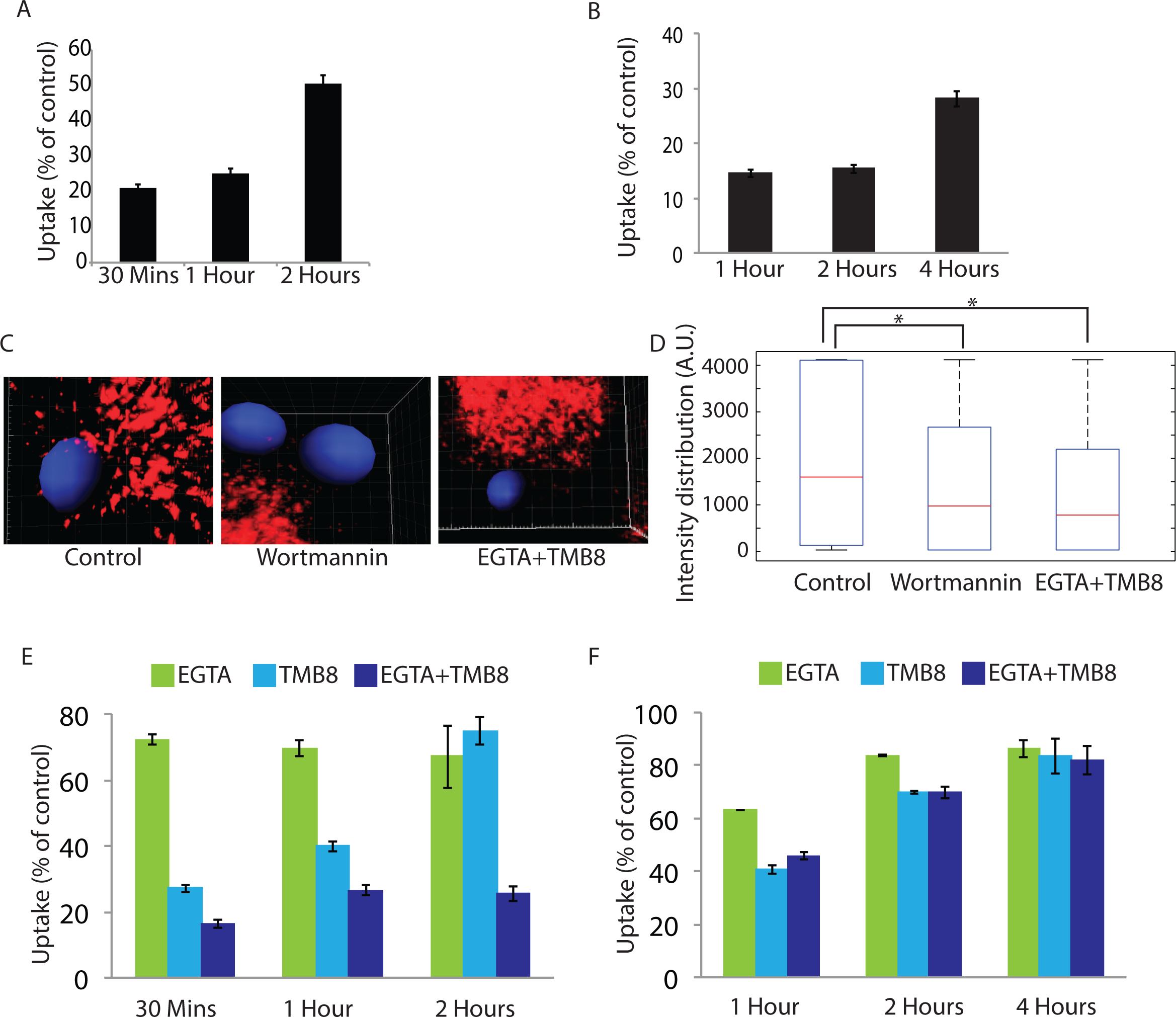
Role of PIP3 and Calcium from intra and extra-cellular sources in recruitment of Mannosidase II vesicles: **A**) THP-1 derived macrophages were pre-treated with 5μM wortmannin for 4 hours. FITC latex beads (1μm) were added to these. The cells were fixed at respective time points and analyzed by flow cytometry. Data shown are average from three different experiments and represented as % uptake in treated cells with respect to the untreated cells (values ± S.D.). B) THP-1 derived macrophages were pre-treated with 5μM wortmannin for 4 hours. They were infected with PKH labeled H37Rv. At respective time points the cells were fixed and analyzed by flow cytomtery. Data shown are average from three different experiments and represented as % uptake in treated cells with respect to the untreated cells (values ± S.D.). C) mCherry-MAN2A (red) expressing RAW264.7 macrophages were pretreated with 5μM Wortmannin or EGTA (3mM)+TMB8 (100 μM) for 30 minutes respectively. The cells were incubated with mouse serum coated latex beads for 30 minutes or 1 hour. At the respective time points, cells were fixed and analyzed by confocal microscopy. The images shown are representative from the 30 minutes samples. The 3D construction of latex bead was done using spot creation module of Imaris 7.2. D) RAW264.7 macrophages expressing mCherry-MAN2A (red) were incubated with mouse serum coated latex beads for 30 minutes and 1 hour. The total intensity of mCherry-MAN2A puncta on the bead surface was determined using 3D spot creation module in Imaris 7.2 and the intensity distribution of the population has been plotted. The box plot represents data from more than 200 beads analyzed from two different experiments (*p-value<0.05). E) THP-1 derived macrophages were pre-treated with EGTA (3mM), TMB8 (100μM) and EGTA+TMB8 for 30 minutes before addition of FITC latex beads (1μm). At respective time points the cells were fixed and analyzed by flow cytometry. Data shown are average from three different experiments and represented as % uptake in treated cells with respect to the untreated cells (values ± S.D.). F) THP-1 derived macrophages were pre-treated with EGTA (3mM), TMB8 (100μM) and EGTA+TMB8 for 30 minutes before addition of PKH labeled H37Rv. At respective time points the cells were fixed and analyzed by flow cytometry. Data shown are average from three different experiments and represented as % uptake in treated cells with respect to the untreated cells (values ± S.D.).

### Extracellular Calcium (Ca^2+^) is needed for phagocytosis and focal exocytosis of Golgi-derived vesicles at the site of phagocytosis

We next wanted to understand the immediate early mediators of focal exocytosis during phagocytosis. Focal exocytosis of vesicles derived from the recycling endosomes as well as lysosomes depends on the functioning of key Ca^2+^ sensors in these compartments (10,11,13). It was therefore imperative to test the role of Ca^2+^ in the focal movement of Golgi-derived vesicles. Release of Ca^2+^ from intracellular stores is one of the key signaling events during the course of phagocytosis and downstream maturation of the phagosomes (35). Expectedly, phagocytosis of both latex beads and H37Rv in THP-1 macrophages was severely hampered in the presence of TMB-8 (Fig. 7E and 7F), an inhibitor of IP3 receptor that serves as the Ca^2+^ release channel from intracellular stores like ER upon binding with IP3 (36). Intriguingly, presence of Ca^2+^ chelator EGTA in the extracellular media also inhibited phagocytic uptake of both latex beads and H37Rv in THP-1 macrophages (Fig. 7E and 7F) thereby suggesting the involvement of extracellular Ca^2+^ in the phagocytic uptake. However it could simply imply the effect of capacitative Ca^2+^ influx, which typically requires the CRAC channels and happens immediately after the intracellular stores are exhausted of Ca^2+^ (36). Treatment with both TMB-8 and EGTA had additive effect on the uptake of latex beads but not on the uptake of *Mtb* by THP-1 macrophages (Fig. 7E and 7F). We also noted a much more pronounced and sustained effect of TMB-8 and EGTA+TMB8 (~70% and ~85% inhibition respectively, Fig. 7E) on the uptake of latex beads as against that of *Mtb*(~25% and ~45% inhibition respectively, Fig. 7F). As observed in the case of wortmannin treatment, inhibition of Ca^2+^ signaling also resulted in a marked decline in the recruitment of Mannosidase-II at the nascent phagosomes (Fig. 7C and 7D).

### Entry of extracellular Ca^2+^ through voltage-gated Ca^2+^ channels helps create the foci for the recruitment of vesicles during phagocytosis

The role and mechanism of recruitment of extracellular Ca^2+^ during phagocytosis is limited to the capacitative influx (37) or through passive accumulation of Ca^2+^ in the phagosomes during phagocytosis (38). An increase in the local Ca^2+^ concentration around phagosomes has been reported to facilitate phagocytosis (39). Interestingly, the role of extracellular Ca^2+^ during membrane repair process and focal exocytosis of vesicles to the damaged site has been extensively reported (40-42). Similarly during neurotransmitter release in the neuronal cells, activation of voltage gated Ca^2+^ channels during action potential helps focal exocytosis of secretory vesicles (43). We hypothesized that one of the earliest signals during phagocytic uptake could consist of Ca^2+^ entry into the cells through the voltage-gated Ca^2+^ channel. In THP-1 macrophages treated with loperamide hydrochloride or amlodipine besylate (inhibitors of L/P-type and L-type voltage gated Ca^2+^ channels respectively) (44,45), phagocytosis of *E. coli* declined considerably (Fig. 8A). These treatments were also effective against phagocytosis of latex beads and *Mtb* in THP-1 macrophages (Fig. 8B and 8C respectively). Treatment with any of these two inhibitors also blocked Mannosidase-II recruitment at the nascent phagosomes during uptake of latex beads in THP-1 macrophages (Fig. 8D). Thus voltage-gated channel appeared to play an important role during phagocytosis of diverse targets.

**Figure 8:**
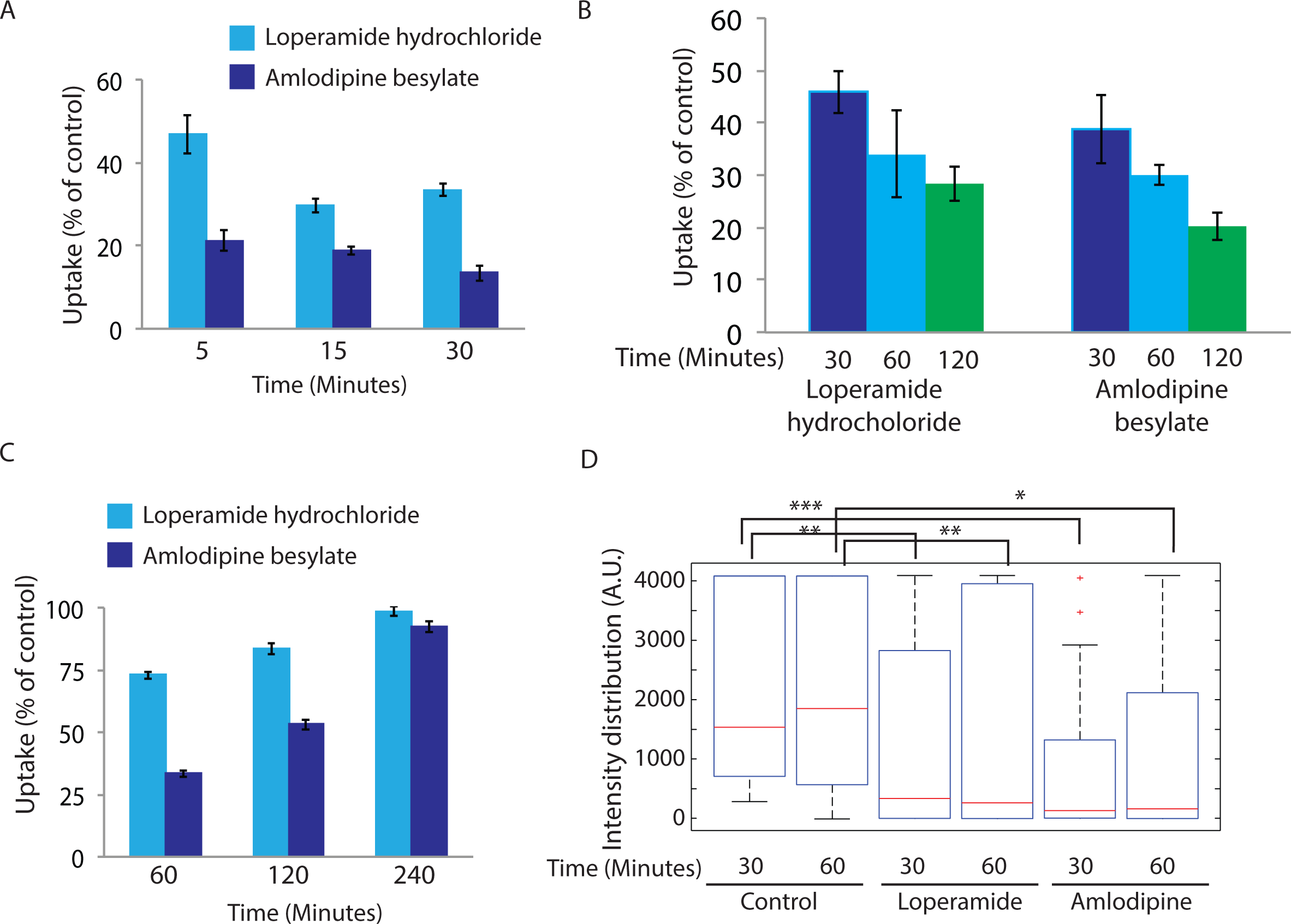
Ca^2+^ entry through voltage gated Ca^2+^ channel helps phagocytosis. **A**) THP-1 derived macrophages were pre-treated with Amlodipine (100μM) or Loperamide (100μM) for 30 minutes before addition of GFP expressing *E. coli*. At respective time points the cells were fixed and analyzed by flow cytometry. Data shown are average from three different experiments and represented as % uptake in the treated cells with respect to the untreated cells (values ± S.D.). **B**) THP1 derived macrophages were pre-treated with loperamide hydrochloride(100μM) and amlodipine besylate(100μM) for 30 minutes and incubated with FITC labeled latex beads. The cells were fixed at respective points and analyzed by flow cytometry (values ± S.D.). **C**) THP1 derived macrophages were pre-treated with loperamide hydrochloride(100μM) and amlodipine besylate(100μM) for 30 minutes and infected with PKH67 labeled H37RV. The cells were fixed at respective points and analyzed by flow cytometry (values ± S.D.). **D**) RAW264.7 macrophages expressing mCherry-MAN2A (red) were pre-treated with amlodipine (100μM) or loperamide (100μM) for 30 minutes and subsequently incubated with mouse serum coated latex beads for 30 minutes and 1 hour. The total intensity of mCherry-MAN2A puncta on the bead surface was determined using 3D spot creation module in Imaris 7.2 and the intensity distribution of the population has been plotted. The box-plot represents data from more than 200 beads analyzed from two different experiments (*p-value<0.05, **p-value<0.01 and ***p-value<0.005).

It now seemed plausible that extracellular Ca^2+^ helped decide the foci for the recruitment of Golgi-derived vesicles during phagosome formation (43). To test that, we investigated the accumulation of PIP3 at the site of engulfment. Inhibition of PIP3 by wortmannin treatment abrogates the uptake as observed in this study (Fig. 7) as well as reported earlier (14). PIP3 is known to interact with the Pleckstrin Homology (PH) domain of proteins. In cells expressing AKT-PH domain fused with mCherry (AKT-PH-mCherry), at 5 minutes post-addition of *E. coli*, we observed, expectedly, a significant accumulation of PIP3 at the site of bacterial entry (Fig. 9A). Also there was a gradual decline in the PIP3 levels as we move further into the cells from the site of phagocytosis (Fig. 9A). However in cells pre-treated with the Ca^2+^ chelator (EGTA, 3mM), the selective accumulation of PIP3 at the site of phagocytosis was lost (Fig. 9A) and PIP3 was almost uniformly distributed across the cell irrespective of the site of engagement with the bacteria (Fig. 9A). Surprisingly, similar effect on PIP3 distribution was observed when these cells were pre-treated with either loperamide or amlodipine (Fig. 9A). In both these cases, the distribution of PIP3 in the cells was not influenced by the site of engagement with the bacteria (Fig. 9A). In cells treated with the PI3K inhibitor wortmannin, AKT-PH-mCherry was as expected, more uniformly distributed (Fig. 9A). Quantitatively, it was evident from the signal intensity plots (see methods) in figure 9B, that a gradient of PIP3 is established during phagocytosis, with the highest concenration at the site of entry. Moreover since blocking the entry of extracellular Ca^2+^ either through chelators or inhibitors of voltage gated channels resulted in a loss of PIP3 gradient in a similar fashion as in the case of wortmannin treatment, it was evident that the formation of PIP3 gradient was dependent on Ca^2+^, more likely on the entry of Ca^2+^ from extracellular milieu through the voltage-gated Ca^2+^ channels (Fig. 9B). Together these results imply that the extracellular Ca^2+^, entering through a voltage-gated Ca^2+^ channel helps forming the foci for the recruitment of PI3K and initiate the signaling cascade for phagocytic uptake in the macrophages.

**Figure 9:**
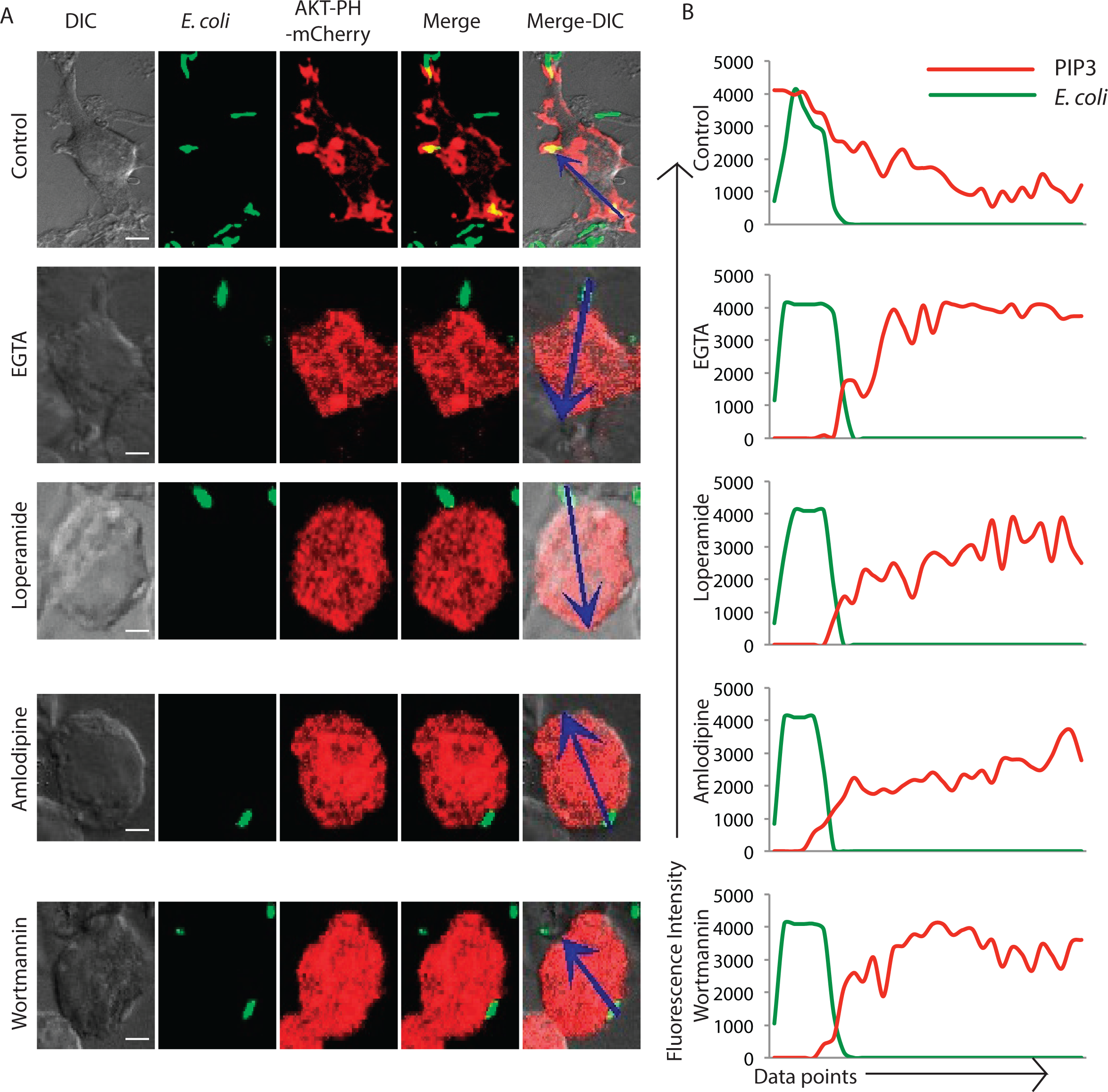
Role of Voltage-gated Ca^2+^ channels in establishing the PIP3 gradient during phagocytosis. **A**) RAW264.7 macrophages were transfected with AKT-PH-mCherry. At 24 hours of cells were incubated with GFP expressing *E. coli* for 5 minutes. In parallel we also had AKT-PH-mCherry expressing cells that were pre-treated with EGTA (3mM), loperamide (100μM), amlodipine (100μM) or wortmannin (5uM) followed by incubation with GFP expressing *E. coli* for 5 minutes. Samples were fixed at 5 minutes and analyzed by confocal microscopy. The arrows in the extreme right image in each of the panel highlight the fluorescence intensity measurements for the analysis presented in figure 9B (scale bar: 4μm). **B**) Images in figure 9A (arrows) were analyzed using intensity profile line tool in the NIS-elements software (see methods). The data represent median from more than 20 fields for each case.

### MAN2A positive vesicles represent Golgi-derived secretory vesicles

During the membrane repair and neurotransmitter release, secretory vesicles derived from the Golgi are rushed to the site of repair or release (41). Fusion of secretory vesicles with the membrane results in closure of membrane damage or release of the vesicular content (41). The fusion involves interaction between v-SNARE VAMP2 on the secretory vesicles and t-SNARE Syntaxin 1 at the plasma membrane (46). We did observe VAMP2 in the phagosome proteome thereby providing a clue that the Mannosidase-II vesicles could most likely be secretory vesicles derived from the Golgi. We therefore tested the uptake of *E. coli* in cells where VAMP2 was knocked down using siRNA. In VAMP2 knockdown cells, *E. coli* uptake was reduced by ~50% at 10 minutes and ~35% by 20 minutes (Fig. 10A). In U937 cells expressing mCherry-MAN2A, we observed upon siRNA treatment recruitment of Mannosidase-2 at the phagosomes declined by nearly 50% at 10 minutes to ~75% at 30 minutes (Fig. 10B). Interestingly, there was no decline in the recruitment of VAMP3 to the *E. coli* phagosomes in VAMP2 knockdown cells (Fig. 10C). That there was progressive loss of Mannosidase-II positive *E. coli* phagosomes in the absence of VAMP2 indicates a compensatory mechanism whereby in the absence of secretory vesicles, membrane from other sources like recycling endosomes, ER etc. get recruited to help phagocytosis. Since VAMP2 is required for the fusion of secretory vesicles with the target membrane, loss of Mannosidase-II at the phagosomes in the absence of VAMP2 suggests inhibition of fusion of Golgi-derived vesicles with the phagosomes. However we could not rule out if VAMP2 knockdown also influenced vesicle release from the GA.

**Figure 10:**
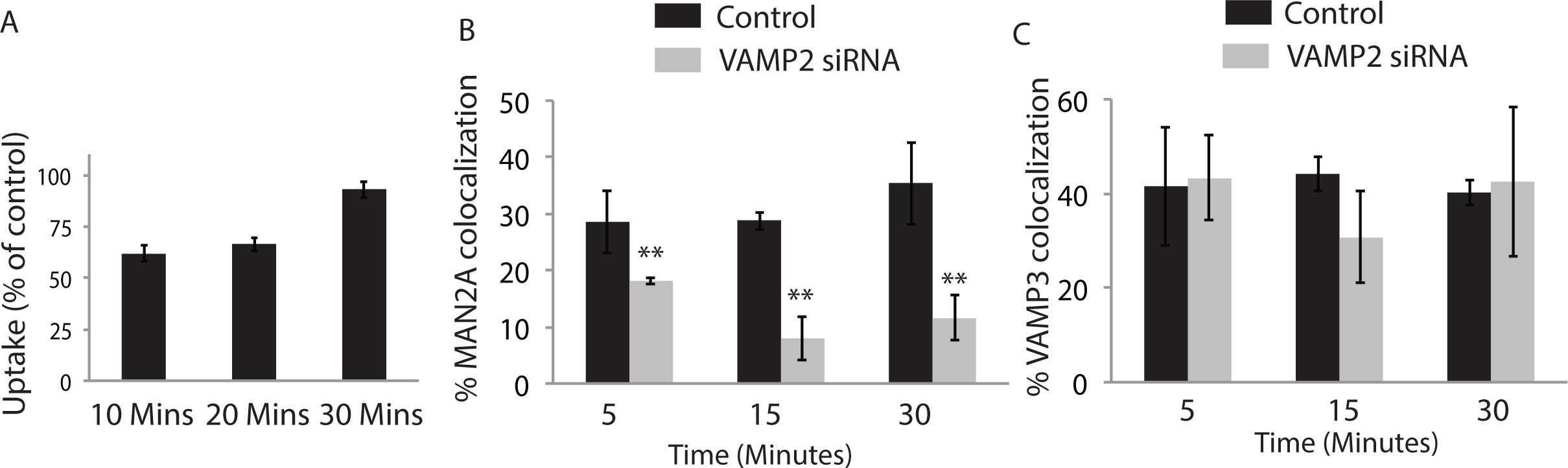
Recruitment of Mannosidase-II at phagosomes require secretory vesicle specific v-SNARE VAMP2. (A) THP1 derived macrophages were transfected with VAMP2 siRNA and at 48 hours post transfection, uptake of *E. coli* was assayed in comparison to the control cells that received scrambled siRNA. Uptake of PKH67 labeled E coli was monitored at 10, 20 and 30 minutes. Data represents average from three replicates and error bars are SD. (B) In mCherry-MAN2Aexpressing cells, VAMP2 was knockdown and incubated with GFP expressing *E. coli* for different time points as mentioned. Samples were then fixed and analyzed through confocal microscope. Data represents average of more than 200 bacteria from two different experiments (values ± S.D.; scale bar). (C) In the experiment mentioned above in B, samples were also stained with anti-VAMP3 antibody followed by Alexa405 labeled secondary antibody and analyzed through confocal microscope. Data represents average of more than 200 bacteria from two different experiments (values ± S.D).

### Mannosidase-IIpositiveearly phagosomes are also NCS1 positive

Having established the role of Ca^2+^ in regulating the movement of Mannosidase-II vesicles during phagocytosis, we next wanted to understand how the focal movement of Golgi-derived vesicles is triggered in phagocytosing macrophages. NCS1 is an EF-hand motif containing protein, originally believed to exclusively express in the neuronal cells (47). Subsequently it was shown to get expressed in a variety of cell lines as TGN resident Ca^2+^ interacting protein (47). NCS1 was reported to be involved in the recruitment of Golgi-derived vesicles during plasma membrane repair in macrophages and was integral to these vesicles (23,48). In cells expressing mCherry-MAN2A we stained for NCS1 during uptake of latex beads or *Mtb*. In both the cases, NCS1 co-localized with the cargo (Fig. 11A and 11B). In the case of *Mtb*, nearly all *Mtb* containing phagosomes were also positive for NCS1 whereas some NCS1 positive phagosomes did not contain Mannosidase-II (Fig. 11B). Similarly for beads, NCS1 intensity was much higher compared to Mannosidase-II intensity (Fig. 11B). Moreover, we observed NCS1 to colocalize with Mannosidase-II in vesicles that were not part of the phagosome (Fig. 11C), corroborating with the fact that both Mannosidase-II and NCS1 are components of the secretory vesicles derived from the GA (23,48).

**Figure 11:**
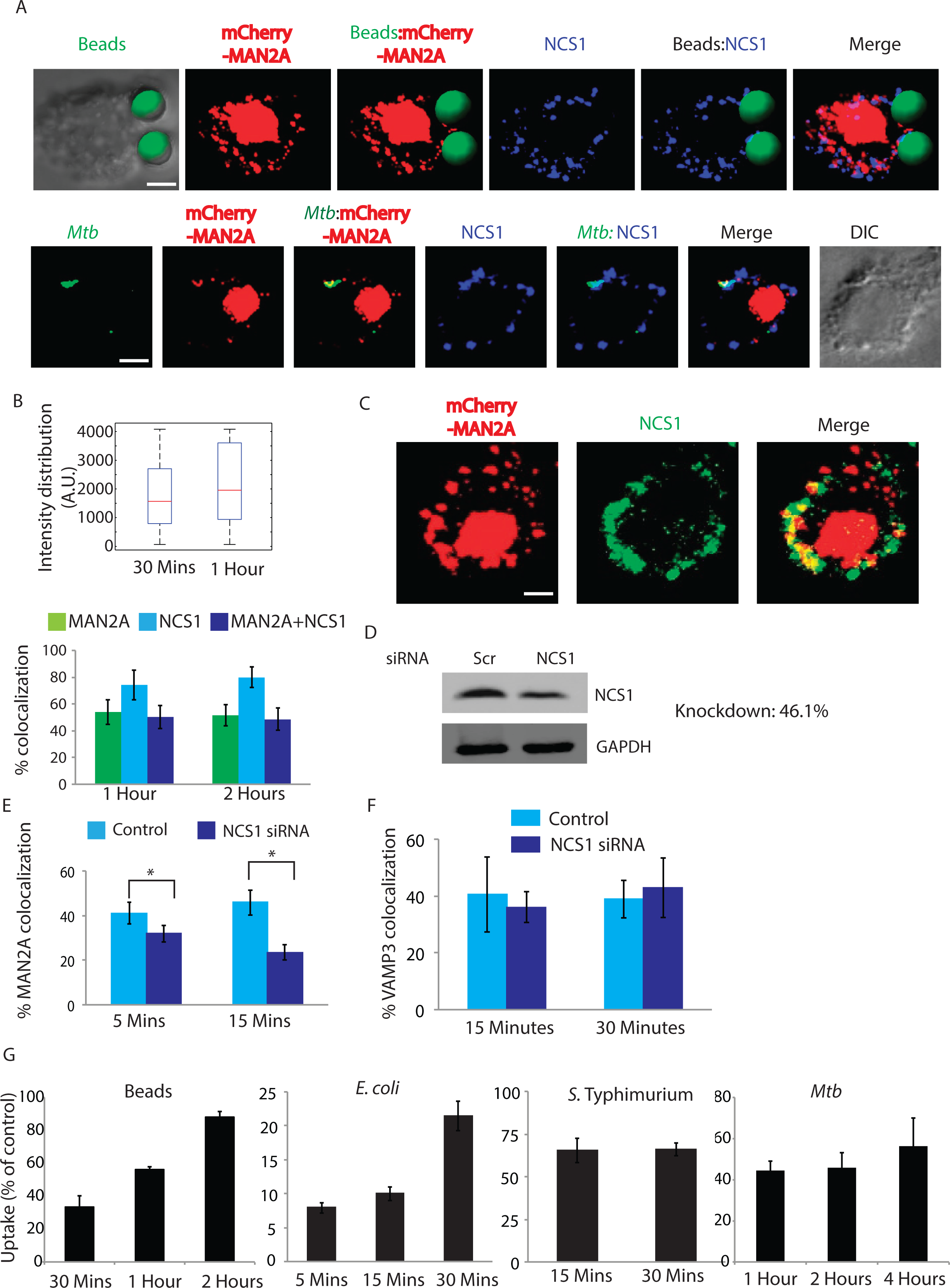
The neuronal calcium sensor (NCS1) in the Golgi apparatus recognizes Ca2+ signal for focal release of Mannosidase-II vesicles. A) mCherrry-MAN2A expressing U937 macrophages were incubated with latex beads (30 minutes) or infected with PKH67 labeled H37Rv (1 hour). Cells were fixed and stained with anti-NCS1 antibody followed by Alexa 405 labeled secondary antibody. For the upper panel latex beads were given green pseudo-color using Imaris. (Scale bar:4μm). B) For the upper panel, in U937 macrophages, incubated with beads for 30 minutes or 1 hour, samples were stained with anti-NCS1 antibody. Presence of NCS1 at the bead surface was calculated using the 3D spot creation module in Imaris 7.2 software. The box-plot at the right shows data from more than 100 beads from two independent experiments. For lower panel, mCherry-MAN2A (red) expressing U937 derived macrophages were infected with PKH67 labeled H37Rv (green) for 1 and 2 hours. At the respective time points, samples were fixed and stained with anti-NCS1 antibody followed by Alexa-405 tagged secondary antibody. The images are representative from the 1hour time point. Percent co-localization of H37Rv with Mannosidase-II, NCS1 or both Mannosidase-II and NCS1 was calculated using Imaris 7.2. The data represents average of more than 150 bacteria from three different experiments (values ± S.D.). C) Co-localization of Mannosidase-II and NCS1 in U937 macrophages. U937 macrophages expressing mCherry-MAN2A were stained with anti-NCS1 antibody followed by secondary antibody (pseudo-colored green). (Scale bar:4μm). D) siRNA mediated knockdown of NCS1 was confirmed by Western blots on the whole cell lysates from the transfected cells. Knockdown was monitored at 48 hours post transfection. E) THP-1 derived macrophages treated with NCS1 siRNA were incubated with GFP expressing *E. coli* for 5 and 15 minutes. Cells were stained with anti-Mannosidase-II antibody to assess the recruitment of Mannosidase-II at the phagosomes in NCS1 depleted cells. Percent co-localization of *E. coli* with Mannosidase-II was calculated using Imaris 7.2. The data represents average of more than 150 bacteria from three different experiments (values ± S.E.M, *p-value<0.05; scale bar: 2μm). F) In the similar experiment as mentioned in E, samples were stained with anti-VAMP3 antibody to assess the recruitment of VAMP3 at the phagosomes in NCS1 depleted cells. Percent co-localization of *E. coli* with Mannosidase-II was calculated using Imaris 7.2. The data represents average of more than 100 bacteria from three different experiments (Value± S.D). G) THP-1 macrophages were treated with siRNA against NCS1 or scrambled control. At 48 hours post siRNA treatment, cells were monitored to uptake latex beads (1μm), *E coli*, *Salmonella* Typhimurium. or H37Rv for indicated time points. Data are shown as % uptake in the siRNA treated cells with respect to the scrambled siRNA control treated cells. Data are representative of three independent experiments (values ± S.D.).

### Depleting Neuronal Calcium Sensor 1 (NCS1) inhibits focal exocytosis of MAN2A vesicle but not of VAMP3 vesicles during phagocytosis

To test whether Ca^2+^ dependent trigger of vesicular trafficking relied on the ability of NCS1 activation upon Ca^2+^ binding, we compared the recruitment of Mannosidase-II at the nascent phagosome during phagocytosis of *E. coli* in THP-1 macrophages that were either treated with scrambled siRNA control or NCS1 specific siRNA (Fig. 11D). Knocking down NCS1 resulted in more than 50% decline in the recruitment of Mannosidase-II at the early *E. coli* phagosomes (Fig. 11E). Requirement of NCS1 was very specific for Mannosidase-II recruitment since in NCS1 depleted cells there was no decline in the recruitment of VAMP3 at the phagosomes (Fig. 11F). Expectedly, knockdown of NCS1 in THP-1 macrophages resulted in a marked decline in the uptake of latex beads, *E. coli*, *Salmonella* and *Mtb* (Fig. 11G). True to all other treatments, which inhibited the phagocytic uptake, NCS1 siRNA knockdown also had both quantitative and kinetic effects on the uptake of all of the targets except in *Salmonella* where the effects were persistent (Fig. 11G). Thus uptake in case of latex beads was ~35, 60 and 85% of the control in the siRNA treated sets at 30 minutes, 1 hours and 2 hours respectively (Fig. 11G). For *E. coli* these numbers were 8, 10 and 22% of control at 5, 15 and 30 minutes (Fig. 11G). In case of *Salmonella*, uptake was about 65% of the control set at 15 and 30 minutes while in case of *Mtb*, the relative uptake in the NCS knockdown cells was 40, 45 and 55% of control at 1 hour, 2 hours and 4 hours respectively (Fig. 11G).

## Discussion

Professional phagocytes like macrophages require continuous supply of membrane in order to form phagosomes around the phagocytosed particles (49). It is now understood that cellular compartments like recycling endosomes, lysosomes and ER could supply membrane for the nascent phagosomes (7,8)(6). Adding to the existing pool of membrane sources available for phagosome formation, in this study we show recruitment of Golgi-derived vesicles at the site of phagocytosis in macrophages. However unlike previous reports, where the studies were mostly restricted to either latex beads or select organism, we show here a more universal requirement of the Golgi-derived vesicles during phagocytosis by macrophages using inert particles, non-pathogenic bacteria and two different pathogenic bacterial species as the cargo. It is important to note here that GA serves as the origin for most vesicles in the cell, including recycling endosomes, endocytic machinery and vesicles destined to the plasma membrane for exocytosis (50). Bacterial pathogens once inside the host cells further subvert the membrane trafficking pathways to ensure prolonged survival and escape from innate defense mechanisms(51). Interestingly GA serves as the hub for trafficking inside the cells (52). It therefore may make strong sense that GA gets involved and alerted of incoming pathogen while the cell has just started to engulf it. This line of investigation seems extremely fascinating at present as it may open a new understanding in the functioning of innate immune system. However involvement of Golgi apparatus during phagocytosis was ruled out earlier (16). Coincidentally all of these reports involved studies on Fcγ-Receptor mediated phagocytosis using IgG-coated latex beads (16). A detailed analysis of uptake of latex beads coated with either IgG or serum revealed the selectivity in the involvement of Golgi apparatus during phagocytosis. While uptake of IgG-coated latex beads was not sensitive to Brefeldin A treatment, uptake of serum opsonized latex beads was sensitive to this treatment. We realized that serum opsonized latex beads would engage a different set of receptors like complement receptor, mannose receptor and scavenger receptors on the macrophages unlike IgG-coated beads, which would only engage with Fcγ receptors(3). Thus while we observed recruitment of Golgi-derived vesicles to most of the cargos studied and that the recruitment of Golgi-derived vesicles was important for their uptake, it was not the case for uptake of IgG-coated beads. Interestingly Brefeldin A treatment also led to a decline in Mannosidase-II recruitment at the nascent phagosomes. More importantly Brefeldin A treatment did not have influence on the recruitment VAMP3, a recycling endosome vesicle specific v-SNARE, which is known to get recruited at early phagosomes. Thus recruitment of Mannosidase-II happened through a pathway other than the established recycling endosome pathway of vesicle recruitment. Better understanding of specific signaling events downstream to these receptors could provide answers on how differential receptor engagement ensures recruitment of selective organelles.

Role of Golgi-derived vesicles in supplying the membrane at plasma membrane is also well known in another context ‐ the membrane repair pathway (23). Using Mannosidase-II as specific marker for Golgi-derived vesicles they showed participation of GA and lysosomes in the repair process through targeted exocytosis (23). The movement of vesicles during phagocytosis from the recycling endosomes and lysosomes was also reported to follow a similar targeted exocytosis where vesicles are directed towards the phagocytic cup (7,13,53). Interestingly more than 90% of Mannosidase-II positive phagosomes at the site of entry were also found to be positive for VAMP-3 and dynamin-2. While VAMP-3 gets recruited through focal exocytosis, dynamin-2 is known to regulate post-Golgi transport of vesicles and facilitate focal exocytosis (13). Dynamins have earlier been implicated in regulating the focal exocytosis in phagocytosing macrophages (13,34,54). Inhibition of dynamins by dynasore treatment resulted in a reduced phagocytic uptake, which may simply occur due to inhibition of dynamin-mediated scission of phagosomes from the membrane (55). However dynasore treatment also resulted in reduced Mannosidase-II recruitment at the nascent phagosomes, which can only happen if dynasore is inhibiting the generation and focal movement of Golgi-derived vesicles. Moreover, we categorically show that Mannosidase-II recruitment is independent of recruitment of markers from endocytic pathways, re-iterating the fact that dynasore mediated effect was mostly due to inhibition of post-Golgi transport of vesicles, consistent with previously reported function of this inhibitor (13). High co-localization of Mannosidase-II with TfR further emphasized very early recruitment of Golgi-derived vesicles. Yet another instance where vesicle exocytosis from Golgi has been extensively studied is in the context of neurotransmitter release, which shows remarkable similarity with the focal exocytosis during membrane repair. Thus it is intriguing to witness the brilliance of cellular economy, where a common mechanism could be utilized to address three entirely independent cellular requirements. Next to understand how target recognition at the cell surface for phagocytic uptake could trigger GA to elicit the movement of the vesicles, we took cues from membrane repair pathway where the damage is typically sensed via entry of extracellular Ca^2+^ into the cells (40,41). We indeed observed inhibiting either extracellular Ca^2+^ or the release of Ca^2+^ from intracellular stores resulted in a loss of phagocytic function in the macrophages. The possibility of a membrane breach during phagocytic uptake in macrophages has never been discussed. Thus most plausible source for the entry of extracellular Ca^2+^ was some membrane channels, which are classically involved at a similar step during neurotransmitter release(43,56). Experiments with loperamide and amlodipine strongly support a critical involvement of voltage-gated Ca^2+^ channel in the extracellular Ca^2+^ entry thereby facilitating phagocytosis. Incidentally the role of voltage-gated channel in regulating podosome formation in the macrophages was recently shown (57). Yet another channel TRPV2 was recently shown to be important for phagocytosis in macrophages and its absence resulted in loss of Ca^2+^ influx from the extracellular milieu and abrogated phagocytic uptake (58). Curiously macrophages lacking TRPV2 were also defective in chemoattractant-evoked motility (58). It may not be unusual to assume that some of the inhibitory effects shown in this study were most likely due to inhibition of TRPV channels at the plasma membrane. Therefore it seems macrophage membrane depolarization could be a more general mechanism of cellular functioning including phagocytosis, adherence and motility.

Phagocytosis was also dependent on the release of Ca^2+^ from intracellular stores, as TMB8 treatment inhibited phagocytosis. A combination of EGTA and TMB8 had more dramatic effects on phagocytosis, which also showed some sort of selectivity in terms of the cargo. Thus, for latex beads, blocking both intra- and extra-cellular cargo abolished their uptake by the macrophages, however in case of bacteria, the block in uptake was more kinetic in nature. It strongly supports the possibility that Ca^2+^ may be extremely critical for the uptake of cargos where cognate receptors are not known/available thereby rely a lot on the membrane depolarization and associated Ca^2+^ entry. The graded importance of Ca^2+^ during phagocytosis of diverse targets needs further exploration for better understanding. Inhibition of Ca^2+^ also resulted in reduced Mannosidase-II recruitment at the phagosomes.

The role of PI3Kinase in phagocytosis is well known, which supposedly helps pseudopod extension while engulfing the target (14). However, it has also been shown that inhibition of PI3Kinase may limit the membrane availability (15). We indeed observed uptake of *Mtb*, *Salmonella* or latex beads by macrophages were severely compromised in the presence of wortmannin. *Salmonella* entry is known to occur via either macropinocytosis or phagocytosis, however the former is insensitive to PI3Kinase inhibition(59). Thus the strains used in this study are taken up by the macrophages through phagocytosis. Interestingly vesicle recruitment has also been reported to be important for membrane ruffle formation, which helps in *Salmonella* macropinocytosis (60). We also noted significantly reduced recruitment of Mannosidase-II vesicles at the nascent phagosomes in the presence of wortmannin. Thus recruitment of Golgi-derived vesicles at the nascent phagosomes indeed required PIP3. Experiments with AKT-PH-mCherry clearly support the establishment of PIP3 gradient during phagocytosis. Selective PIP3 enrichment has previously been shown regulating cellular polarity, chemotaxis and pseudopod extension (14,61,62). The most dramatic observation was however the use of voltage-gated Ca^2+^ channels by macrophages to set the foci for PIP3 accumulation, which eventually results in all the downstream signaling and recruitments. Interestingly, dynamins have conserved PH- domain at their C-terminal, making them responsive to the PIP3 levels. It has been shown that mutations in the PH domain of dynamin could result in severe defects in its key functioning like post-Golgi transport and endocytosis (63,64). Thus PI3K and dynamins seem to work together for the focal recruitment of Golgi-derived vesicles during phagosome biogenesis.

The Ca^2+^ sensing function in the GA is attributed to the resident molecule NCS1, which was initially discovered in the neuronal cells regulating the synaptic transmission (47). NCS1 was also shown as part of the secretory vesicles (65). In macrophages, ablation of NCS1 resulted in a block in the membrane-resealing pathway (23). We therefore hypothesized; NCS1 could also serve as the Ca^2+^ sensor in Golgi apparatus for triggering the exodus of the vesicles during phagocytosis. Interestingly, NCS1 knockdown not only reduced the phagocytic function of macrophages, there was also significantly lower recruitment of Mannosidase-II at the phagosomes. Further emphasizing on the existence of a separate mechanism for Mannosidase-II recruitment, NCS1 knock down did not influence recruitment of VAMP3 on the early phagosomes.

The results form purified latex beads phagosomes supported the overall observation in this study that phagocytic uptake is a complex process and involves several players. We deliberately focused on the molecules below 25kDa cut-off to make sure enrichment of three key classes of molecules including RABs (~25 kDa), ARFs (~20 kDa) and VAMPs (~14 kDa). This was to ensure that the presence of other high abundance and high molecular weight proteins did not mask these low abundant low molecular weight proteins. We identified 17 different RABs on the phagosomes out of which nine were associated with Golgi function. We also identified five out of six known ARFs, where ARF1 and ARF3 were particularly important for their known association with Golgi apparatus, phagosome/engulfment and exocytosis(66-68). We could identify four different VAMPs in the phagosomes including VAMP2, 3, 4 and 8. While role of VAMP3 in phagosome biogenesis is well described, VAMP2 is particularly known for its involvement in exocytosis, neurotransmitter release and secretory vesicles (28,29). Previously in a proteomic study of the latex-beads containing phagosome, VAMP4 was identified as phagosome membrane associated protein (69). VAMP4 are trans-Golgi resident proteins and are also involved in the immature secretory granules (70). We also found SEC22B in the phagosome MS data, an important SNARE known to be important for the recruitment of ER during phagocytic uptake (16). Considering the number of RABs, ARFs and VAMPs associated with the exocytosis and secretory vesicles, it seems very likely that the true identity of the Golgi-derived vesicles that are recruited during phagosome biogenesis could be secretory vesicles.

At the molecular level it seems SNARE mediated fusion may be involved in the phagosome biogenesis. We were able to track the presence of syntaxin 1, the t-SNARE at the plasma membrane involved in SNARE mediated vesicle fusion, along with Mannosidase-II at the early phagosomes (data not shown). Syntaxin-1 is known to facilitate fusion of secretory vesicles(76). Moreover, its presence will also be required for the fusion of VAMP3 positive vesicles, a vSNARE typically located in the recycling endosomes. However presence of VAMP2 in the proteomic data, a secretory vesicle specific v-SNARE, gave us a clue that possibly recruitment and fusion of Mannosidase-II vesicles to the phagosomes happen through the secretory pathway. By knocking down VAMP2, we were able to see selective decline in the recruitment for Mannosidase-II without effecting VAMP3 recruitment at the phagosomes. VAMP2 knockdown also resulted in a marked decline in the uptake, reiterating the fact that phagocytosis involves cooperation between multiple sources, which can provide membrane for membrane biogenesis. Taken together the results in this study allowed us to reconstruct the process of phagocytosis (Fig. 12). It is important to note that Golgi-derived vesicles are one of the several sub-cellular vesicular pools available for phagosome biogenesis including recycling endosomes and lysosomes. It actually explains why in many cases except for PI3K inhibition, the effects were more kinetic in nature.

**Figure 12:**
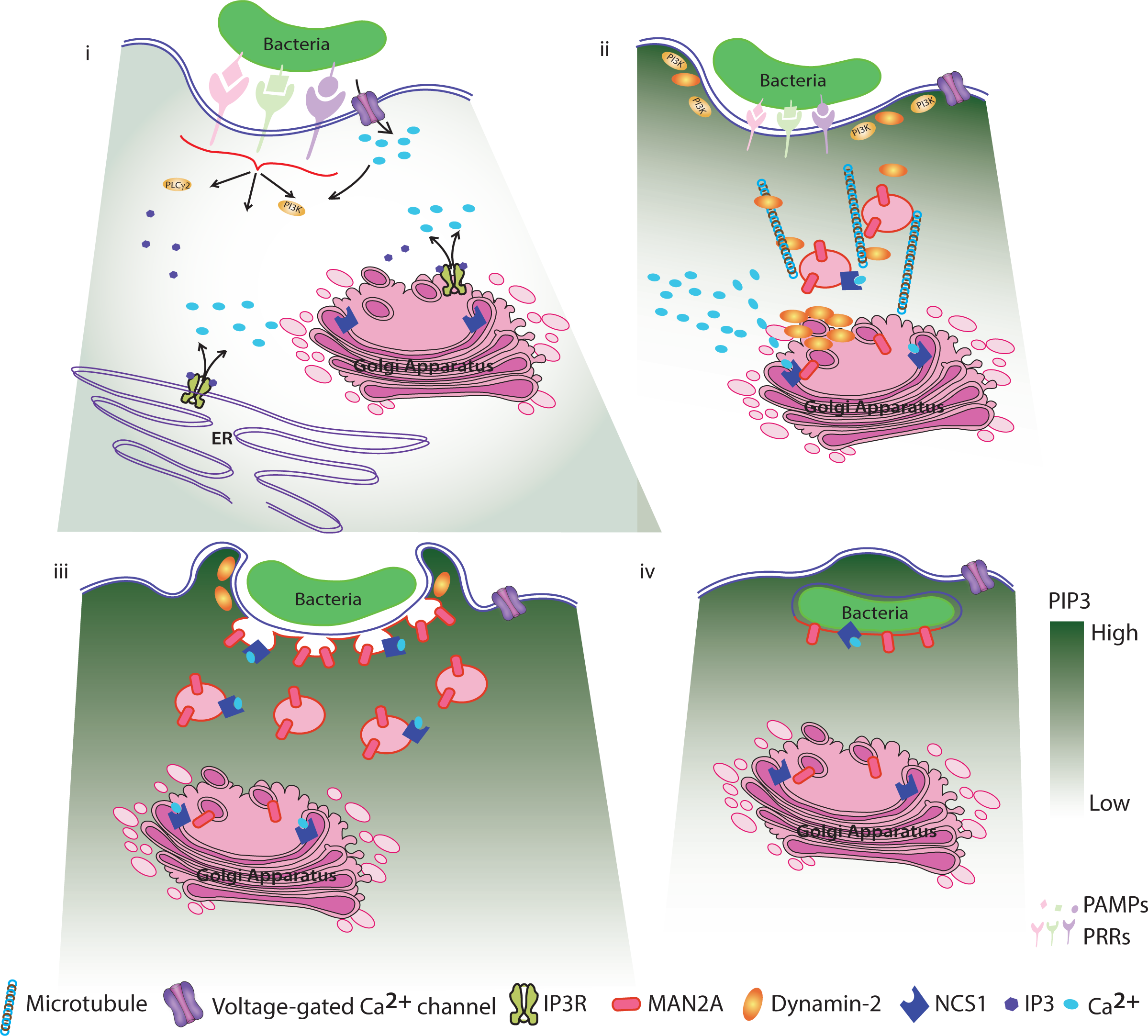
A schematic model describing involvement of Golgi-derived vesicles during phagocytic uptake in macrophages. i) Initial recognition of an object (bacteria, beads, cell debris etc) for phagocytosis results in membrane depolarization possibly due to torsional stress and resulting in the activation of voltage gated Ca^2+^ channels, leading to the entry of extracellular Ca^2+^ into the cells. The entry of Ca2^+^ through voltage gated Ca^2+^ channels sets the focus for quick and efficient recruitment of PI3K and results in the accumulation of PIP3 at the site of phagocytosis. These early events are further aided by signaling through specific pattern recognition receptors and release of Ca^2+^ from intracellular stores. ii) Increased cytosolic Ca^2+^ is sensed by Golgi-resident NCS1, which triggers the movement of Mannosidase-II vesicles towards the site of phagocytosis, guided by a gradient of PIP3, along with dynamin and microtubule. iii) Mannosidase-II vesicles fuse with the membrane at the site of phagocytosis and help grow the phagosome around the cargo before final scission and internalization. iv) The membrane of the nascent phagosome is contributed in part by the plasma membrane and rest from the Golgi-derived vesicles (in this model for clarity, we have excluded lysosome and recycling endosomes as additional sources, see text).

In conclusion, we show here an as yet unknown function of Golgi-derived vesicles during phagocytic uptake in macrophages. The targeted exocytosis coupled with phagocytosis has been studied in the past, however involvement of secretory vesicles from the GA during phagocytosis was unprecedented. Thus while the utilization of distinct phagocytic pathways by pathogens to escape killing in macrophages is well known, this study highlights the redundancy in the repertoire of membrane sources recruited downstream to distinct phagocytic receptors. How the membrane repertoire utilized during phagocytosis influences the fate of infection poses an attractive direction of investigation. Finally our finding that the voltage-gated Ca^2+^ channel could play a role in the process of phagocytosis may be potentially harnessed in future for developing better therapeutic interventions for various infectious diseases including tuberculosis.

## Methods

### Ethics statement

The animal experiments were performed upon prior approval from the institutional ethics committee (IAEC) of International Center for Genetic Engineering and Biotechnology (Approval no.: ICGEB/AH/2013/03/IMM-38).

#### Reagents And Antibodies

The following reagents were used in this study: Wortmannin (Sigma Aldrich,W1628), Dynasore Hydrate (Sigma Aldrich, D7693), Nocodazole (Sigma Aldrich, M1404), TMB8 (Sigma Aldrich, T111), EGTA (Amresco, 0732), Loperamide (Sigma Aldrich, L4762), Amlodipine besylate (Sigma Aldrich, A5605), PKH 67 (Sigma Aldrich, MINI 26), RIPA buffer (Amresco,N653), BSA (Sigma Aldrich, A2153), Saponin (Sigma Aldrich, 47036), Puromycin (Invivogen, ant-pr-1), 4 μm aldehydate-sulphatelatex beads (Life Technologies, A37304) and 1μm yellow green aldehyde latex beads (Life Technologies, F8823). All the siRNA used in this study were siGenome siRNA SmartPool (Dharmacon Inc). The transfection reagent used for siRNA transfections was Dharmafect-2 (Dharmacon Inc). The primary antibodies used in this study are: Mannosidase II (abcam, ab12277), Dynamin (Santacruz, sc-6401), VAMP3 (abcam, ab43080), Transferrin (abcam, ab84036). The secondary antibodies used in this study are: Alexa fluor 405 and Alexa fluor 568 conjugates from (Life Technologies).

#### Plasmid Constructs

The plasmid constructs used in this study are mCherry-ManII-N-10 (Addgene plasmid #55073), pcDNA3.1_AktPH-mCherry (Addgene plasmid #67301), pCT‐ Golgi‐ GFP (CYTO104-VA-1) and pCT-Mem-GFP (System Biosciences, CYTO100-PA-1).

#### Bacterial Culture Maintainence

Mycobacterial cultures were maintained in 7H9 media (Difco) supplemented with 10% OADC. Single cell suspension of Mycobacterial strains were prepared by aspiration of the culture eight times with 26 gauge needle and six times 30 gauge needles. Quantification of this prepared culture was done by taking absorbance at 600 nm wavelength (0.6 O.D. corresponds to ~100 x 10^6^ bacteria). The bacteria thus appropriately calculated were added to the cells at the mentioned MOI (Multiplicity of Infection). For microscopy experiments, the desired number of bacteria was stained with PKH67 (Sigma Aldrich), a lipophilic green fluorescent dye, as per the manufacturer’s protocol. The stained bacteria were then passed thrice through a 26-gauge needle and used for infection. *E*. *coli* and *Salmonella* Typhimurium strains were maintained in LB (Difco). For infection culture density was determined by OD at 600 for both *E. coli* and *S*.Typhimurium containing GFP, an OD of 1= 1 x 10^8^ bacteria for both these strains.

#### Tissue Culture

THP-1 cells (a kind gift from Dr. Dong An, UCLA) and U937 cells (ATCC) were cultured in RPMI 1640 medium (Life Technologies) and RAW264.7 (ATCC) murine macrophages were grown in Dulbecco’s modified Eagle’s media (DMEM, Life Technologies) supplemented with 10%Fetal Bovine Serum (FBS, GIBCO) and maintained 37°C in a humidified, 5% CO_2_ atmosphere. THP1 cells were differentiated with 20ng/μl PMA for 24 hours, washed with plain RPMI and maintained in 10% FCS supplemented RPMI for another 24 hours. The cells were then infected with respective bacteria/beads. RAW264.7 macrophages were seeded in respective plates and differentiated with 200ng/ml LPS for 12 hours. The cells were washed once with plain DMEM and infected with respective bacteria/beads.

#### Latex Bead Experiments

Latex beads were coated with anti-human IgG(for THP1 and U937 macrophages) or anti-mouse IgG (for RAW264.7 macrophages) as described by (16), 1mg/ml antibody for 1 hour or incubated with 10 times the volume of mouse/human sera for 1 hour and then utilized for further experimentation by confocal or flow cytometry.

#### Animals and isolation of BMDMs

Bone marrow derived macrophages (BMDMs) were isolated from femurs of BALB/C mice (4-6 weeks old, female) obtained from institutional animal house. BMDMs were obtained by culturing the marrow cells in the presence of macrophage-colony stimulating factor (M-CSF, eBioscience, 14-8983-80) for 7 days. Fully differentiated macrophages were harvested and seeded for infection with H37Rv. The infection protocol was same as described above for THP-1 macrophages.

#### Experiments with Latex beads

For microscopy experiments 4μm aldehyde sulfate latex beads (Life Technologies) were incubated with human/mouse IgG on an agitator overnight. The beads were washed twice with plain DMEM by centrifugation at 2000 rpm for 10 minutes and added to cells at respective MOI. For flow cytometry experiments, 1μm FITC labeled aldehyde beads were added at respective MOI to the differentiated THP1 cells.

#### Transfection & Nucleofection Assays

Transfection & Transduction was carried out using JetPrime Reagent (Himedia) as per the manufacturer’s protocol. Nucleofection was carried out with the 4D-Nucleofector™ System Lonza in 20 μl Nucleocuvette™ Strips with program DS-136, as per the manufacturer’s protocol. At 24 hours post Nucleofection these cells expressed the inserted vector as determined by visualization with a fluorescence microscope.

#### siRNA Transfection & Inhibitor Assays

Post 24 hours of PMA treatment, the cells were washed once with plain RPMI. The siRNA’s were added as per the manufacturer’s protocol. Post 48 hours of incubation the respective infection/assays were performed. For the inhibitor assays such as, Dynasore(40μM, 80μM), Wortmannin (as specified) were added 4 hours before infection. While Nocodazole(25 μM) was added 2.5 hours before infection and EGTA(3mM), TMB8(100 μM), Loperamide hydrochloride(100 μM) and Amlodipine besylate(100 μM) were added for 30 minutes before infection.

#### Creation of stable cell lines

The stable cell lines were created from lentivirus cytotracer plasmids (B4GALT1-mRuby and NEUM-EGFP, System Biosciences). 1X106 U937 cells were added to a 24 well plate. The harvested media containing lentivirus (refer to transfection section) was added to them. At 72 hours post infection the cells were selected on Puromycin at 350ng/ml for 21 days. The population of positive cells was routinely checked by microscopy and it was found that post selection for 21 days the population of cells remained stable (in our case 30-40%).

#### Staining for Confocal Microscopy

At specific time points cells were fixed with 4% Paraformaldehyde (PFA) for 15 minutes. This was followed by 2 washes with 1x PBS. The cells were permeabilized with 0.4% Triton X 100 for 30 minutes and washed once with 1X PBS. Blocking was performed using 3 %( w/v) BSA and 0.5%Tween-20 in 1X PBS for one hour, followed by one wash with 1X PBS. The cells were now stained with respective primary antibody made in the blocking solution at specific dilution for 90 minutes. The coverslips were then washed once with 1X PBST and twice with 1X PBS. This was followed by staining with respective secondary antibody tagged with fluorophore of choice for 90 minutes. The primary and secondary antibodies were diluted in blocking solutions for use. After three washes with 1X PBS, the coverslips were then mounted on glass slides with anti-fade reagent(Life Technologies). Images were acquired with Nikon A1R Laser Scanning Confocal Microscope with a 100X/1.4NA Plan Apochromat VC, DIC N2 objective lens. Image processing viz. 3D reconstruction, co-localization and intensity measurements were done via Imaris 7.2 (Bitplane).

#### Live Cell Imaging

The live cell imaging dish pre-seeded was RAW264.7 macrophages expressing mCherry-MAN2A. The live cell system was set at 5% CO2 and 370C. The field was set and imaged for 10 minutes. This was followed by an on-stage addition of 0.0001% saponin for 5 minutes. The cells were washed once, supplemented with complete media and imagining continued. The entire procedure was performed on stage to analyze the pre and post treatment effects on the membrane under our experimental set up.

#### Flow Cytometry experiments

At respective time points, the infected cells were washed thrice with sterile 1X PBS to remove extracellular bacteria/beads and fixed with 2% PFA. The cells were scraped and run on BD FACS Canto or BD FACS Influx cytometer’s.

#### Western Blot

Post SDS PAGE, the proteins were blotted onto a nitrocellulose membrane using a semidry-transfer system. Following incubation with primary and secondary antibodies, the blots were scanned using Odyssey InfraRed Imaging System (LI-COR BioSciences, Lincoln, NE, USA) at various intensities in order to obtain a blot scan with minimum background. All settings were rigorously maintained for all experiments. The scans were quantitatively analyzed using Odyssey InfraRed Imaging System Application Software_Version.3.O (commercially available from (LI-COR BioSciences, Lincoln, NE, USA).

#### Phagosome Isolation

Latex bead phagosome for 1μm latex beads were isolated as per protocol described earlier (77). At 1 hour post addition, the plates were washed 4 times with cold 1XPBS, ensuring all the extracellular beads were completely washed off. The cells were scraped off with a rubber policeman and centrifuged twice with cold 1XPBS at 1200 RPM for 5 minutes. The cells were pooled and washed once with homogenization buffer at 1200 RPM for 5 Minutes (3mM imidazole, 250 mM sucrose, pH 7.4). The cells were then incubated in appropriate volume of homogenization buffer (with protease inhibitor cocktail – Amresco) for 30 minutes. The cells were lysed by dounce homogenizer till 90% of the cells were disrupted as observed under a light microscope. This lysate was centrifuged at 1200 RPM for 5 minutes at 40°C to remove the unbroken cells. This was followed by preparation of sucrose gradient for ultracentrifugation. The homogenate was made to be consisting of 40% sucrose by mixing with equal volume of 62% sucrose and 2.25 ml of the same was layered upon 2.0ml of 62% sucrose solution. We then added 2.25 ml each of 35%, 25% and 10% sucrose solutions. The tubes were centrifuged at 100,000g for 1 hour at 40C in an SW28 rotor (Beckman Coulter). The LBC band obtained between 10 and 25% sucrose layers was collected and washed once with cold 1XPBS at 40,000g at 40C in an SW28 rotor (Beckman Coulter).

#### Mass Spectrometry & Data Analysis

For LC-LTQ Orbitrap MS analysis, samples were re-solubilized in 2% [v/v] acetonitrile, 0.1% [v/v] formic acid in water and injected onto the trap column at a flow rate of 20 μl/min subsequently peptides were separated on Zorbax 300SB-C18 (Agilent, Santa Clara, CA, USA) by a gradient developed from 2% [v/v] acetonitrile, 0.1% [v/v] formic acid to 80% [v/v] acetonitrile, 0.1% [v/v] formic acid in water over 180 min at a flow rate of 300 nl/min onto an Agilent 1200 (Agilent, Santa Clara, CA, USA) nano-flow LC-System that was in line coupled to the nano-electrospray source of a LTQ-Orbitrap discovery hybrid mass spectrometer (Thermo Scientific, San Jose, CA, USA). Full MS in a mass range between m/z 300-2,000 was performed in the Orbitrap mass analyzer with a resolution of 30,000 at m/z 400 and an AGC target of 2x 105. The strongest five signals were selected for CID (collision induced dissociation)-MS/MS in the LTQ ion trap at a normalized collision energy of 35% using an AGC target of 1x 105 and two microscans. Dynamic exclusion was enabled with one repeat counts during 45 s and an exclusion period of 180 s. Peptide identification was performed by CID-based MS/MS of the selected precursors. For protein/peptide identification, MS/MS data were searched against the *Mus musculus* amino acid sequence database (downloaded in August 2015) using an in-house Mascot server (version 2.4) through the Proteome Discoverer 1.4 software. The search was set up for full tryptic peptides with a maximum of three missed cleavage sites carbamidomethyl on cysteine, and oxidized methionine were included as variable modifications. The precursor mass tolerance threshold was 10 ppm, and the maximum fragment mass error was 0.8 Da. The significance threshold of the ion score was calculated based on a false discovery rate of <1%, estimated by the peptide validator node of the Proteome Discoverer software.

#### Analysis for functional classes

The non-redundant list of proteins identified was matched against a series of gene ontology classes from AMIGO2 database. The selection of gene ontology was empirically made based on known and perceived classes, which together could represent the set of trafficking proteins identified. The obtained matches were then represented as a network using Cytoscape 3.2.0.

#### Statistical Analysis

Comparative groups were analyzed using paired two-tailed t-test using inbuilt function in MS Excel.

#### 3-D recreation and analysis

The intensity of a particular fluorophore on the bead was estimated by first creating a three dimensional bead surface in the captured image (z-stack) using “spot module” in Imaris Version 7.2 (Bitplane) which automatically detects spheres in an image depending upon the dimensions fed. The software then allows one to determine fluorescence intensity on the surface individually for each fluorophore. For each case maximum fluorescence intensity was determined and plotted.

#### PIP3 gradient analysis

The gradient of fluorescence in the AKT-pH mCherry experiments was determined using “intensity profile line tool” in NIS-Elements (NIKON).

## Acknowledgements

The work was supported by a grant from the Department of Biotechnology (DBT), Govt. of India (BT/PR14730/BRB/10/874/2010, DK) and partly by International AIDS Society and CFAR funded CNIHR grant from NIH, USA (DK). The work related to *Mycobacterium tuberculosis* infections were carried out at DBT funded TACF facility at ICGEB. We thank Prof. Sarman Singh for access to the flow-cytometer at AIIMS and Purnima for her help in the confocal microscopy. We thank C-CAMP, NCBS, Bangalore for the mass spectrometry facility. NV is a recipient of a senior research fellowship from the University Grants Commission, India.

## Author Contribution

DK and NV designed the study and wrote the paper. NV, SBAB and SSG performed the experiments. NV and DK analyzed the data. MS provided important reagent for the study. All authors reviewed and approved the final version of the manuscript.

